# Copper trafficking in *Trypanosoma cruzi*: the transcriptional response of candidates to balance toxicity and recruitment

**DOI:** 10.1101/2024.04.12.589278

**Authors:** Marcelo L. Merli, María G. Mediavilla, Xinyu Zhu, Paul A. Cobine, Julia A. Cricco

## Abstract

*Trypanosoma cruzi* (Chagas disease) depends on acquiring nutrients and cofactors, like copper (Cu), from its hosts. Cu is essential for aerobic organisms, but it can also be toxic, so its transport and storage must be regulated. In the present study, we characterized the effects of changes in Cu availability on growth, intracellular ion content, and oxygen consumption. Our results show that Cu is essential for epimastigote proliferation and for metacyclogenesis, while intracellular amastigotes suffered from Cu stress during infection. We identify several genes potentially involved in Cu metabolism among which orthologs of the conserved P-type Cu ATPases involved in Cu export and loading of secreted enzymes were found and named *Tc*CuATPase. *Tc*CuATPase transcription is regulated during infective stages and by Cu availability in epimastigotes. No homologs were identified for the high affinity importer CTR1 instead we propose that the iron transport *Tc*IT a ZIP family transporter is involved in Cu uptake based on its transcriptional response to Cu. Further canonical Cu targets (based on homology to yeast and mammals) such as the iron reductase *Tc*FR and the cupro-oxidase *Tc*Fet3 are up regulated during infective stages and under intracellular Cu stress. We also demonstrated that Cu, iron, and heme metabolisms are related. In sum, Cu metabolism is essential in *T. cruzi* life cycle. Even though cytosolic Cu-chaperons are still missing, we propose a model for Cu transport and intracellular distribution in *T. cruzi* including conserved factors such as *Tc*CuATPase and others such as *Tc*FR and *Tc*IT playing novel functions.

## Introduction

Copper (Cu) is an essential ion for organisms of all kingdoms, especially for aerobic organisms. The redox potential of the Cu^2+^/Cu^+^ couple is biologically relevant and allows cupro-proteins to catalyze redox reactions. Example of cupro-proteins are cytochrome *c* oxidase-type *aa*3 (a terminal oxidase of the respiratory chain), plastocyanin (an electron transfer center of photosynthesis), NADH dehydrogenase-2 (a bacterial electron transfer enzyme), nitrite reductase, Cu/Zn superoxide dismutase (for superoxide radical detoxification), and different oxidases as laccases or monooxygenases [1,2]. Free Cu ions has the potential to inappropriately bind and disrupt essential iron-sulfur proteins or to cause oxidative damage via Fenton-like reaction [1], so the transport and storage of Cu must be regulated by different importers, exporters, and chaperones involved in its homeostasis. In some organisms, such as yeast, the concentration of cytosolic free copper is near zero because it is efficiently sequestered into organelles as Golgi network, vacuoles, or mitochondria. Copper also forms inert complexes with proteins such as metallothioneins or with the tripeptide glutathione [3].

Chagas disease caused by *Trypanosoma cruzi* is the most prevalent parasitic disease in many countries of America [4]. This protozoan parasite is a trypanosomatid, a group of kinetoplastid among which *Trypanosoma brucei* cause the sleeping sickness disease and *Leishmania* spp. cause different types of leishmaniasis. *T. cruzi* has a complex life cycle characterized by at least four well defined stages displayed in the insect vector and the mammal host: epimastigotes (replicative, non-infective stage), metacyclic trypomastigotes (non-replicative, infective stage), amastigotes (replicative stage), and bloodstream trypomastigotes (non-replicative, infective stage) [5,6]. Each stage is morphologically and metabolically different and depends on acquiring several nutrients and cofactors from the host. *T. cruzi* depends on copper for virulence as previous studies showed that the COX (cytochrome *c* oxidase-type *aa*3) is essential for replication and infectivity of this parasite [7]. *T. cruzi* must then acquire Cu from the different hosts that it encounters along its life cycle through the action of importers and chaperones.

Membranes form the most significant protection against Cu toxicity limiting uptake into the cell and subsequently into organelles. Therefore, identifying the transporters responsible for recruiting Cu is critical to understanding the homeostasis of this element. In mammals Cu can be imported by CTR1 (copper transporter 1) proteins as Cu^+^ ion after its reduction by a Cu^2+^ reductase or taken up as Cu^2+^ by DMT-1 (divalent metal transporter 1) [8]. During Cu excess Cu P-type ATPase is responsible for its export out of the cell. These Cu P-type ATPases also localize to the trans Golgi network (TGN) for delivery of Cu to exported copper proteins or to intracellular vesicles for Cu storage [9]. The Cu P-type ATPases of *Trypanosoma congolense* and *T. brucei* have been experimentally analyzed [10] and the function and expression profile of Cu P-type ATPases of *Leishmania major* was studied during the life cycle of this parasite [11]. In addition, bioinformatic analysis revealed potential P-type ATPases for Cu, Ca, Na, and other ions in other *Leishmania* spp., *T. brucei*, and *T. cruzi* but the function of these proteins has not been experimentally verified [12].

To clarify the copper homeostasis network in *T. cruzi*, in the present study we identified several proteins that could be involved in Cu reduction, import, and distribution in this parasite. The expression of these genes was studied across the life cycle stages of *T. cruzi*: epimastigote, trypomastigote and intracellular amastigote forms. We characterized the effects of changes in Cu ion availability on growth behavior, intracellular Cu content, metacyclogenesis, oxygen consumption, infections, and gene expression for epimastigotes and for amastigotes. Our results show that Cu is an essential ion for epimastigote proliferation and for metacyclogenesis process. We also observed that epimastigotes tolerated high concentrations of Cu, but this caused a negative effect on intracellular amastigote replication. Intriguingly we observed a relationship between Cu, iron, and heme metabolisms and found that Cu can partially stimulate growth of epimastigotes under limited heme and cause a slightly negative effect on iron uptake. This study unveils the conserved factors of *T. cruzi* Cu homeostasis and reveals an apparent lack of some of the Cu importers and intracellular chaperones found in other organisms. Based on the results presented here, we propose a model for Cu transport and intracellular distribution in *T. cruzi*.

## Results

### *Trypanosoma cruzi* presents a non-conserved network for Cu uptake and distribution

We searched for proteins involved in Cu homeostasis in the TriTrypDB database using polypeptide sequences of Cu transporters, chaperones, and cupro-proteins of other organisms (Table 1). The proteins identified in the genome of *T. cruzi* Dm28c strain are summarized in Table 1. We found several copies of a Cu^2+^ reductase (96% of similarity between them) previously described as a Fe^3+^ reductase (*Tc*FR) in *T. cruzi* [13] that is homologue to the Fe^3+^ reductases Fre1 to Fre7 of *S. cerevisiae*. Some of these *S. cerevisiae* Fe^3+^ reductases are reported also as Cu^2+^ reductases [14]. We did not identify homologues to the yeast plasma membrane high-affinity Cu transporters Ctr1, Ctr2, or Ctr3 [15] or to the human Divalent Metal Transporter (DMT-1) [16], but we did find a transporter in *T. cruzi,* previously described as a Fe^2+^ transporter (*Tc*IT), that belongs to the ZIP family [17]. It is worth mentioning that some ZIP proteins can transport divalent cations, such as Cu^2+^ [18,19]. Based on the lack of other candidates we propose this transporter could be a Cu importer.

**Table 1.** Genes identified in *T. cruzi* genome putatively involved in copper homeostasis.

|  | Seed | gene accession number | Description | Identified gene in *T. cruzi* | gene accession number<sup>a</sup> | Localization<sup>b</sup> |
| --- | --- | --- | --- | --- | --- | --- |
| **Plasma Membrane** | Ctr1, Ctr2, and Ctr3 (*Saccharomyces cerevisiae*) | YPR124W, YHR175W and YLR411W | High-affinity Cu transport proteins | - |  |  |
|  | DMT-1 or Nramp2 (*Homo sapiens*) | BAA24933 | Divalent metal transporter, mainly for Fe<sup>2+</sup> but also for Mn<sup>2+</sup>, Cu<sup>2+</sup>, Ni<sup>2+</sup> and Zn<sup>2+</sup> | - |  |  |
|  | Fre1 to Fre7 (*S. cerevisiae*) | YLR214W, YKL220C, YOR381W, YNR060W, YOR384W, YLL051C, and YOL152W | Fe<sup>3+</sup> and Cu<sup>2+</sup> Reductase | TcFR | BCY84_13742, BCY84_13743, BCY84_05177, BCY84_05181, BCY84_05183, BCY84_05184 | cytoplasm (points) |
|  | LIT1 and LIT2 (*Leishmania major*) | LmjF.31.3060 and LmjF.31.3070 | Fe<sup>2+</sup> Transporter. ZIP Family Transporter | TcIT | BCY84_01266, BCY84_15905 | pellicular membrane (weak) |
| **Cytosol** | Atx1 (*S. cerevisiae*) | YNL259C | Cytosolic Cu Chaperone, transports Cu<sup>+</sup> to the secretory vesicle copper transporter Ccc2 | - |  |  |
|  | Cup1-1, Cup1-2 and Crs5 (*S. cerevisiae*) | YHR053C, YHR055C, and YOR031W | Metallothionein; binds Cu<sup>+</sup> and mediates resistance to high concentrations of Cu and cadmium | - |  |  |
|  | Ccs1 (*S. cerevisiae*) | YMR038C | Cu chaperone for superoxide dismutase Sod1 | - |  |  |
|  | Sod1 (*S. cerevisiae*) | YJR104C | Cytosolic copper-zinc superoxide dismutase; detoxifies superoxide | - |  |  |
| **Golgi/ER** | Ccc2 (*S. cerevisiae*) | YDR270W | Cu P-type ATPase, deliver Cu<sup>+</sup> in Golgi network, vesicles, and/or plasma membrane | TcCuATPase | BCY84_03133, BCY84_03134, BCY84_03135 | acidocalcisomes |
|  | Fet3 (*S. cerevisiae*) | YMR058W | Multicopper oxidase involved in Fe<sup>2+</sup> oxidation. It works with Ftr1 transporter in the Fe<sup>3+</sup> transport | TcFet3 | BCY84_15293 | cytoplasm (weak, points, reticulated) |
| Mitochondria | Pic2 (*S. cerevisiae*) | YER053C | Cu and Phosphate mitochondrial Transporter | *TcPic2* | BCY84_14401 | mitochondrion |
|  |  |  |  | *TcPic2*-like | BCY84_10693 | endocytic/cytoplasm |
|  | Sco1 (*S. cerevisiae*) | YBR037C | Cu Chaperone, deliver Cu<sup>+</sup> to the cytochrome c oxidase | *TcSco1* | BCY84_03573, BCY84_12119 | mitochondrion |
|  |  |  |  | *TcSco*-like | BCY84_01670 | mitochondrion (50%) |
|  | Sco2 (*S. cerevisiae*) | YBR024W | Cu Chaperone, deliver Cu<sup>+</sup> to the cytochrome c oxidase | *TcSco2* | BCY84_19910 | (no data) |
|  | Cox11 (*S. cerevisiae*) | YPL132W | Cu Chaperone, deliver Cu<sup>+</sup> to the cytochrome c oxidase | *TcCox11* | BCY84_00471 | Cytoplasm, flagellar cytoplasm, nuclear lumen |
|  | Cox17 (*S. cerevisiae*) | YLL009C | Cu Chaperone, deliver Cu<sup>+</sup> to Sco1 and Cox11 | *TcCox17* | BCY84_15315 | Cytoplasm, flagellar cytoplasm, nuclear lumen |
|  |  |  |  | *TcCox17*-like | BCY84_02381 | cytoplasm (reticulated) |
|  | Cox19 (*S. cerevisiae*) | YLL018C | Chaperone, deliver Cu<sup>+</sup> to the cytochrome c oxidase | *TcCox19* | BCY84_18510 | cytoplasm (reticulated) |
|  | Cox1 (*S. cerevisiae*) | Q0045 | cytochrome c oxidase subunit 1 | *TcCox1* | AKT44498, AKT44499 | (no data) |
|  | Cox2 (*S. cerevisiae*) | Q0250 | cytochrome c oxidase subunit 2 | *TcCox2* | P98023 | (no data) |

<sup>a</sup> Gene accession numbers belonging to *T. cruzi* Dm28c strain 2017 (assembly GCA\_002219105) genome from TriTrypDB, except for TcCox1 and TcCox2 that are NCBI data base gene accession numbers.
<sup>b</sup> Experimental data obtained from TrypTag database, localization of the homologues in *T. brucei*. For the mitochondrial proteins, only C-terminal tagging is considered.

Cu export is mediated by Cu P-type ATPases which are conserved proteins across prokaryotic and eukaryotic organisms [20]. We identified three copies of a gene that encodes for a protein with homology to a Cu P-type ATPase, presenting 99.5 % of similarity between them, and we named it as *Tc*CuATPase. *Tc*CuATPase presents the motifs conserved in P-type ATPases for Cu [12] and could be involved in exporting Cu from the cell at the plasma membrane and/or delivering Cu to cupro-proteins in Golgi network and vesicles. One class of proteins loaded in the Golgi network by the Cu P-type ATPases are the multicopper oxidases. A cupro-protein *Tc*Fet3 was identified as a homologue to the yeast multicopper oxidase Fet3 that oxidizes Fe^2+^ to Fe^3+^ for subsequent uptake by transmembrane transporter Ftr1. Although no homologue to yeast Ftr1 was found in *T. cruzi* genome, the multicopper oxidases could have other functions as, for example, laccase enzymes that oxidize organic compounds such as bilirubin or ascorbate [21]. *Tc*Fet3 presents all the metal binding residues (His81, His83, His126, His128, His413, His416, His418, His483, Cys484, His485, and His489) and two of the three residues involved in ferroxidase specificity (D278 and Y354, but not E185) identified in *S. cerevisiae* Fet3 [22].

Finally, mitochondrial Cu transport seems to be conserved in *T. cruzi*. We found several genes in the *T. cruzi* genome that encode for proteins with homology sequence to the yeast Sco1, Sco2, Cox11, Cox17, Cox19, and Pic2 [1,23]. They are listed in Table 1 and identified as *Tc*Sco1, *Tc*Sco2, *Tc*Cox11, *Tc*Cox17, *Tc*Cox19, and *Tc*Pic2. In yeast, all these proteins are involved in delivering Cu to COX of the respiratory chain. *Tc*Pic2 was named after its *S. cerevisiae* homologue Pic2, but not after the related phosphate carrier Mir1, because *T. cruzi* gene presents the conserved residues specific of the sequence of the copper carrier Pic2 (Cys21, Cys44, and Cys225) [24]. The bioinformatic search also revealed other *T. cruzi* proteins with Pic2-like, Cox17-like, and Sco1/2-like domains, that we identified in Table 1 as *Tc*Pic2- like, *Tc*Cox17-like, and *Tc*Sco-like. *Tc*Sco-like (as *Tc*Sco1 and *Tc*Sco2) presents the conserved residues for Cu binding of Sco chaperones of *S. cerevisiae* (Cys148, Cys152, and His239 of Sco1 [25]). *Tc*Cox17-like and *Tc*Sco-like were conserved among trypanosomatids but not in other non-related eukaryotic organisms.

The subcellular localization of the *T. brucei* homolog proteins was evaluated by TrypTag database (Table 1). The database shows that some of these proteins were detected to the predicted localization sites: *Tb*IT localized to the pellicular membrane, *Tb*CuATPase localized to specialized vesicles of the parasite (acidocalcisomes), and *Tb*Pic2, *Tb*Sco1 and *Tb*Sco-like localized to mitochondria, supporting the hypothetical function we assigned to these proteins. Other *T. brucei* homologues, *Tb*FR, *Tb*Fet3, *Tb*Cox11, *Tb*Cox17, *Tb*Cox17-like, and *Tb*Cox19 (weak or reticular cytoplasmic localization), presented a different cellular localization than expected based on the proposed function.

As mentioned, several proteins from the Golgi network and from mitochondria were identified, but the proteins from the other cellular localizations were not found with the criteria used. At the plasma membrane, no orthologues to the high-affinity copper transporters Ctr1, Ctr2, or Ctr3 were found. No homolog proteins to Atx1 or Ccs1 that would be involved in cytosolic delivery of copper were identified either. We hypothesize that the molecule trypanothione (TrySH), analogue to glutathione (GSH) in trypanosomatids[26], could be involved in Cu cytosolic distribution and storage as GSH was described in other models [15], or that cytosolic chaperone protein/s must be still determined.

### Infective stages have enhanced expression of *Tc*FR, *Tc*IT, *Tc*CuATPase, and *Tc*Fet3

Based on our bioinformatic research, *T. cruzi* does not have the canonically studied copper importers localized in plasma membrane but does have *Tc*FR, *Tc*IT, *Tc*CuATPase, and *Tc*Fet3, potentially localized to the plasma membrane and the Golgi network. We measured the levels of these mRNAs across the *T. cruzi* life cycle: epimastigotes, metacyclic trypomastigotes, amastigotes and cell-derived trypomastigotes (Fig. 1). The expression of the four genes was at least 2-fold higher for both trypomastigote and amastigotes (the infective stages of the parasite) compared to epimastigote stage. *Tc*FR and *Tc*CuATPase genes presented similar patterns: a 2-fold change in metacyclic trypomastigotes and amastigotes, and 3-4-fold change in cell-derived trypomastigotes compared to the epimastigote stage. *Tc*IT and *Tc*Fet3 showed a higher fold change in expression (8 to 13-fold change respect to epimastigotes) in some of the stages: 8-13-fold for *Tc*IT gene in amastigotes and cell-derived trypomastigotes stages, and 13-fold for *Tc*Fet3 gene in metacyclic trypomastigotes. These results suggest that the four genes are important in the infective stages of the parasites.

**Figure 1.**
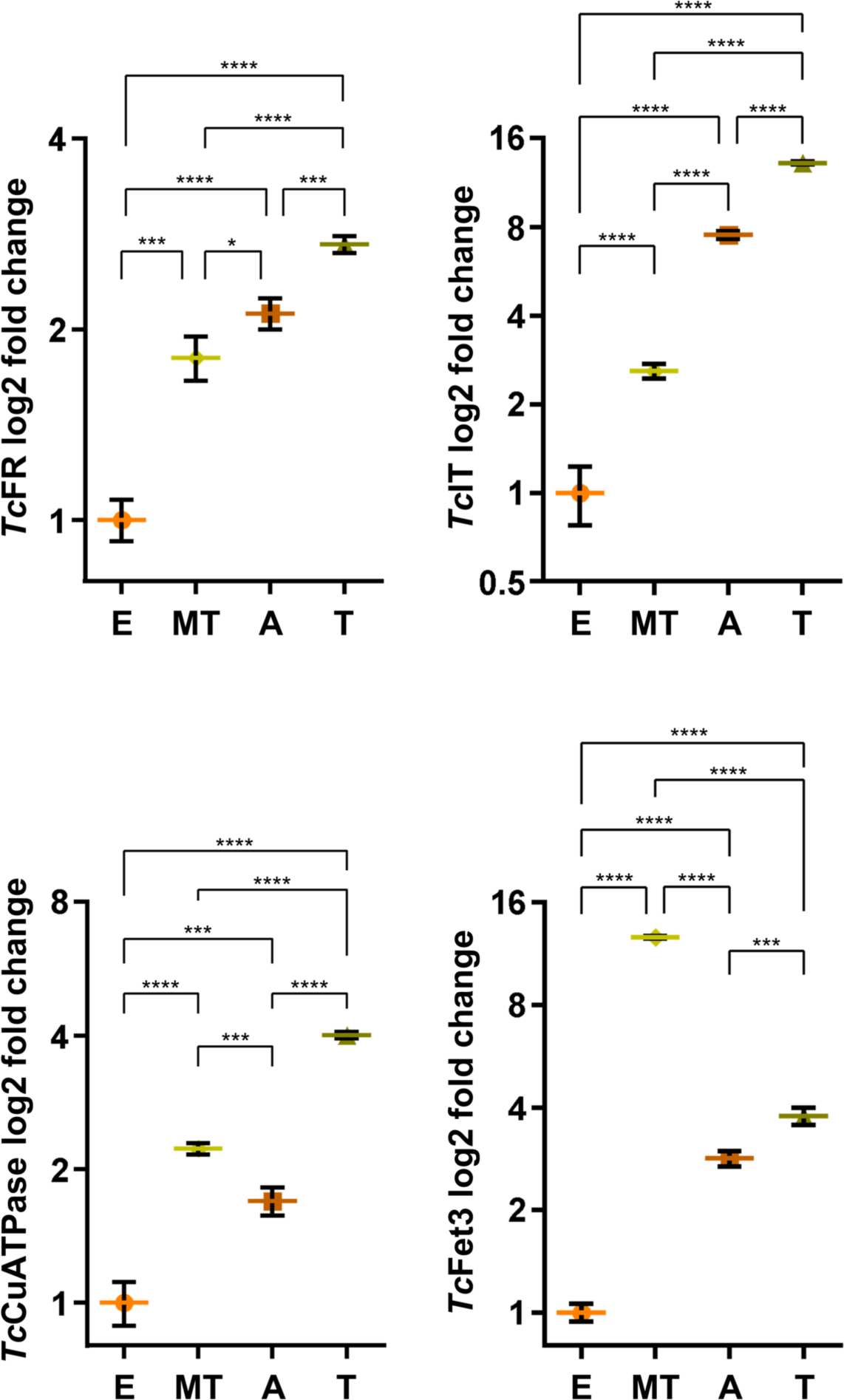
*Tc*FR, *Tc*IT, *Tc*CuATPase, and *Tc*Fet3 genes are overexpressed in the infective stages. Relative mRNA levels of *Tc*FR, *Tc*IT, *Tc*CuATPase, and *Tc*Fet3 genes in the different stages of *T. cruzi* – epimastigotes (E), metacyclic trypomastigotes (MT), amastigotes (A), and cell-derived trypomastigotes (T). The figure shows the log_2_ of fold change of the genes that were determined by 2^-ΔΔCt^ method. *Tc*Ubiquitin was used as housekeeping gene for normalization. Epimastigote stage mRNA content was used as reference (fold change = 1). The results are representative of at least two independent experiments in triplicate. Data represent averages ± SD. Differences were determined using one-way ANOVA and Bonferroni’s multiple comparisons post-hoc test. Significant differences are shown as asterisks: *p <0.05, ***p <0.001, and ****p <0.0001.

### Epimastigotes of *Trypanosoma cruzi* can withstand significant changes in Cu availability

To study the relevance of Cu homeostasis in *T. cruzi* we analyzed the impact of changes in the metal ion availability in the epimastigote stage. Parasites incubated in LIT-10 % FBS-5 μM hemin were challenged to copper stresses for 6 days. High Cu concentrations were obtained by the addition of CuSO_4_ while restrictive conditions were set by the addition of two different copper chelators: bathocuproine disulfonate (BCS, a non-permeable Cu chelator) and neocuproine (NeoCup, a membrane permeable Cu^+^ chelator). Fig. 2 shows that the growth profile of epimastigotes was not severely affected by increasing concentrations of Cu between 100-250 μM but a negative effect was observed at 500 μM (Fig. 2A). The addition of BCS caused a mild negative effect on epimastigote growth, with a reduction of parasite number observed at the stationary phase (Fig. 2B). While the addition of NeoCup severely affected epimastigote growth (Fig. 2C). At the end of the treatment (6^th^ day), the intracellular Cu concentration was measured by ICP-OES. Fig. 2D-E show that epimastigotes incubated with the higher concentrations of CuSO_4_ increased intracellular Cu content while those incubated with BCS presented significantly lower Cu content as chelator concentrations increased, indicating that the treatment applied effectively modulates cellular Cu loading. Instead, parasites treated with NeoCup did not decrease Cu intracellular content (Fig. 2F) but presented a severe growth defect. For the following studies we selected conditions that allowed the parasite survival throughout the assays time span, to measure the response to these treatments.

**Figure 2.**
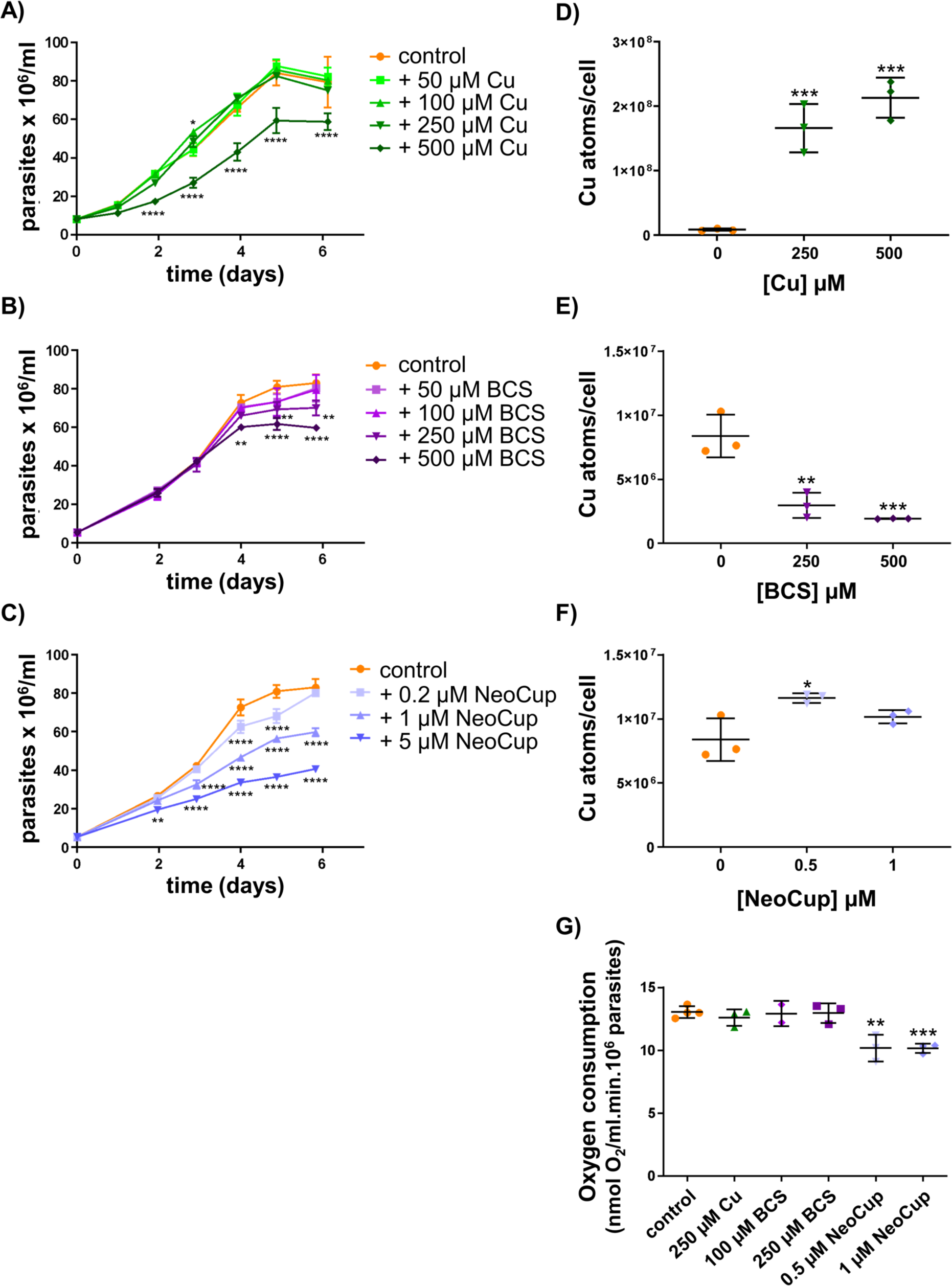
High concentration of copper or its chelation reduces the replication and affects Cu intracellular content of epimastigotes. A-C) Growth curves of *T. cruzi* epimastigote in LIT-10 % FBS-5 μM hemin medium plus different concentrations of copper (Cu), different concentrations of the chelator bathocuproine disulfonate (BCS), different concentrations of the chelator neocuproine (NeoCup), or not addition (control condition). The results are representative of at least two independent experiments. Growth curve data represent averages ± SD. Results were statistically analyzed by two-way ANOVA - Tukey’s multiple comparisons test. Differences compared to controls are marked with asterisks: *p <0.05, **p <0.01, ***p <0.001 and ****p <0.0001. D-F) Cu content determined by ICP-OES of epimastigotes grown for 6 days in LIT-10 % FBS-5 μM hemin medium plus different concentrations of Cu, different concentrations of the chelator BCS, and different concentrations of the chelator NeoCup. The results are representative of at least two independent experiments in triplicate. Data represent averages ± SD. Differences were determined using one-way ANOVA and Dunnett post-hoc test compare with controls. Significances are shown as asterisks: *p <0.05, **p <0.01, and ***p <0.001. G) Oxygen consumption was measured in epimastigotes grown in LIT-10 % FBS-5 μM hemin medium plus Cu, BCS, NeoCup or no addition (control), for 3 days. The results are representative of at least two independent experiments in triplicate. Data represent averages ± SD. Differences were determined using one-way ANOVA and Dunnett post-hoc test compare with control. Significance is shown as asterisks: **p <0.01 and ***p <0.001.

To test mitochondrial function, we measured oxygen consumption of the cells treated with Cu and the chelators for 3 days (exponential growth phase) as is shown in Fig. 2G. Only 0.5 and 1 μM NeoCup treatments significantly decreased the oxygen consumption (22 %); however, no difference was observed between the two concentrations used, suggesting that 0.5 μM of NeoCup was enough to obtain the maximal effect. Interestingly, BCS did not affect oxygen consumption. Then, we repeated this measure at stationary phase (5-day treatment), and no significative differences between treated and control parasites was observed (10.4 ± 0.2 nmol O2/mL.min.10^6^ parasites for 250 µM BCS treatment *vs.* 9.8 ± 0.9 nmol O2/mL.min.10^6^ parasites for the control). These results suggest that the growth defect observed for BCS treatment is not a consequence of the oxygen consumption capacity of the cells.

### Copper, iron and heme homeostasis are connected in *Trypanosoma cruzi*

Copper is required for the multicopper oxidases that are associated with Fe uptake. To analyze the relationship of Cu with iron and heme metabolisms, we first measured the Fe content of cells treated with Cu or Cu chelator for 6 days. The addition of Cu reduced the Fe content (Fig. 3A), but the addition of the chelators did not modify it (Fig. 3B-C). Moreover, the addition of Cu did not modify intracellular heme concentration (Fig. 3D). We also measured the effect on cellular Cu when the heme or Fe contents of the medium were modified (Fig. 4A-B). Treatments with 20 µM hemin or 100 µM Fe (without hemin) reduced epimastigote intracellular Cu while increasing its Fe content, confirming that cofactors are incorporated.

**Figure 3.**
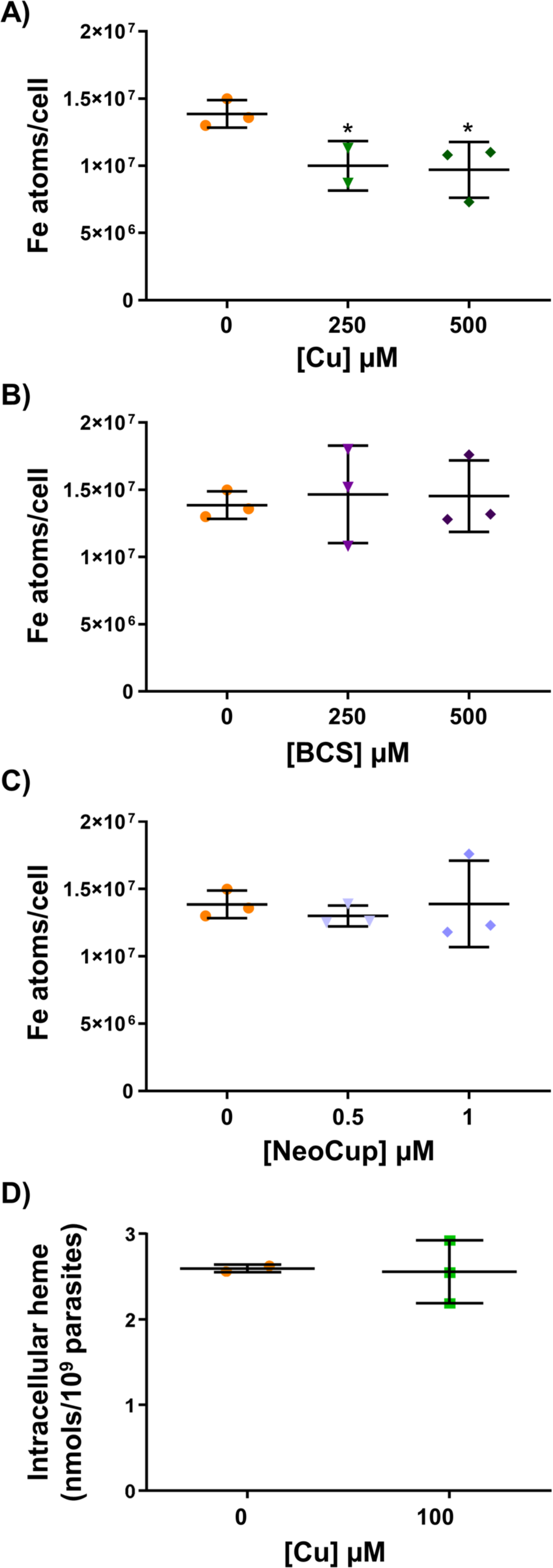
Copper reduces Iron intracellular content in epimastigotes. A-C) Iron (Fe) content determined by ICP-OES of epimastigotes grown for 6 days in LIT-10 % FBS-5 μM hemin medium plus different concentrations of copper (Cu), different concentrations of the chelator bathocuproine disulfonate (BCS), and different concentrations of the chelator neocuproine (NeoCup). The results are representative of at least two independent experiments in triplicate. Data represent averages ± SD. Differences were determined using one-way ANOVA and Dunnett post-hoc test compare with control. Significance is shown as asterisk: *p <0.1. D) Intracellular heme content determined by pyridine method in epimastigotes grown in LIT-10 % FBS-5 μM hemin medium plus 100 μM Cu or without addition (control) for 4 days. The results are representative of at least two independent experiments in triplicate. Data represent averages ± SD. No differences were found using Unpaired t test.

**Figure 4.**
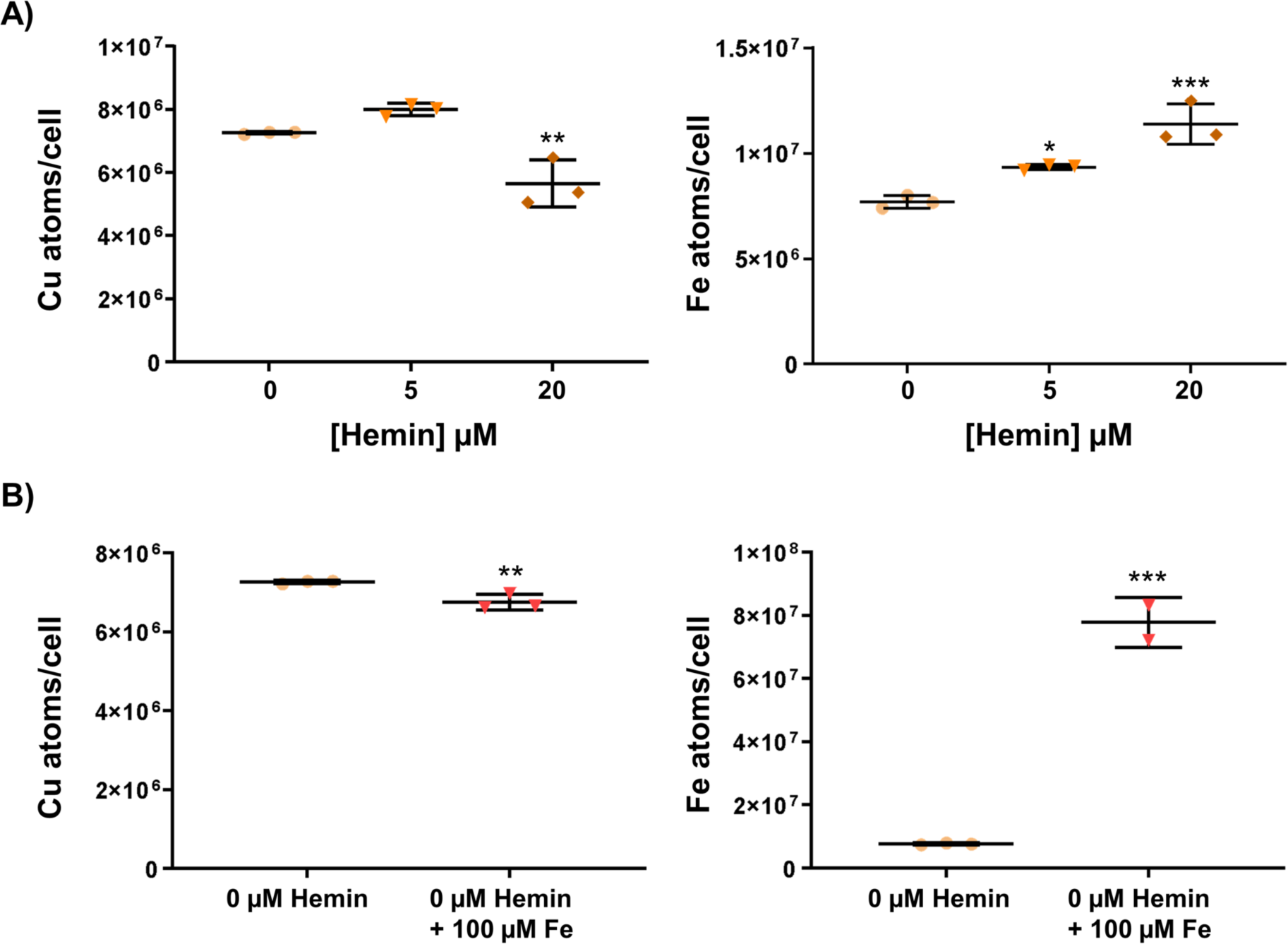
Hemin and iron reduce copper intracellular content in epimastigotes. Parasites were maintained 10 days in LIT-10 % FBS without hemin medium always kept at mid-log phase by periodic dilutions after which the experiments reported in this figure were performed. A) copper (Cu) and iron (Fe) contents were determined by ICP-OES in epimastigotes grown for 6 days in LIT-10 % FBS without hemin medium or plus 5 or 20 µM hemin. Differences were determined using one-way ANOVA and Dunnett post-hoc test compare with control. Significance is shown as asterisks: **p <0.01. B) Cu and Fe contents were determined by ICP-OES in epimastigotes grown for 6 days in LIT-10 % FBS without hemin medium with or without 100 μM Fe. Differences were determined using two-tailed Unpaired t test. Significance is shown as asterisks: **p< 0.01. All the results are representative of at least two independent experiments in triplicate. Data represent averages ± SD.

Then, we tested the effect of Cu addition on hemin starved epimastigotes. Epimastigotes were cultured in LIT medium (LIT-10 % FBS) without addition of hemin for at least 10 days before the assays. Then the cultures were diluted in the same fresh medium supplemented with 5 mM hemin (control) and with different concentrations of Cu, without the addition of hemin. Fig. 5A shows that, as expected, replenishment of 5 μM hemin improved the growth of the epimastigotes when compared with those kept without hemin. Surprisingly, the addition of 50, 100 or 250 μM Cu induced a growth recovery, after 4-5 days of treatment. The number of cells reached similar amount that control parasites, suggesting that Cu could mimic same effect of hemin addition, at least partially until heme storage is completely depleted. As heme can stimulate epimastigote proliferation mediated by the production of a ROS burst, we decided to assay ROS production by Cu [1]. Epimastigotes were cultured without hemin and with or without supplemental Cu. Fig. 5B shows that the addition of Cu significantly increased the production of ROS, specifically superoxide anion, compared to the control. These results suggest that Cu can stimulate the growth of epimastigotes, in absence of extra hemin, by triggering the production of ROS.

**Figure 5.**
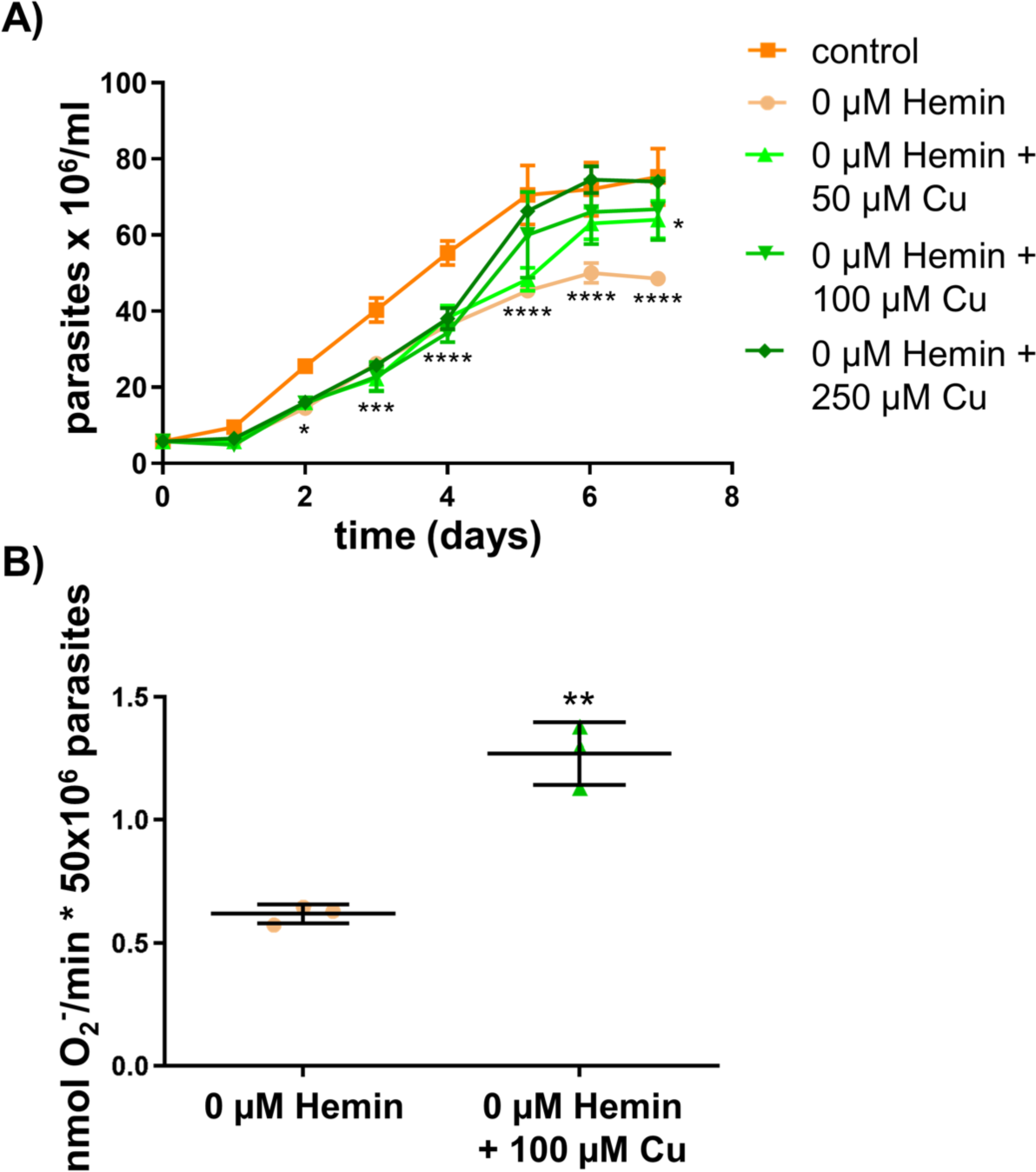
Copper induces epimastigotes growth via ROS production. Parasites were maintained 10 days in LIT-10 % FBS without hemin medium always kept at mid-log phase by periodic dilutions after which the experiments reported in this figure were performed. A) Growth curves of *T. cruzi* epimastigotes in LIT-10 % FBS without hemin (0 µM Hemin) medium plus different concentrations of copper (Cu). Control was performed by adding 5 µM hemin to the same medium (without copper). Two-way ANOVA - Tukey’s multiple comparisons test compares with control. *p <0.05, **p <0.01, ***p <0.001 and ****p <0.0001. B) Superoxide anion production in epimastigotes maintained for 5 days in LIT-10 % FBS without hemin medium plus 100 µM Cu or no addition. Differences were determined using two-tailed Unpaired t test. Significance is shown as asterisks: **p< 0.01. All the results are representative of at least two independent experiments in triplicate. Data represent averages ± SD.

### *Tc*IT and *Tc*CuATPase gene expression respond to Cu-stress

Changes in mRNA level of *Tc*IT, *Tc*FR, *Tc*CuATPase and *Tc*Fet3 genes in response to Cu excess or deficiency (or its chelation) in the epimastigotes were analyzed and are shown in Fig. 6. The treatment with 1 μM NeoCup, which results in intracellular Cu chelation but not in its diminution, caused an increment in mRNA transcripts of all four genes (3 to 4-fold changes). Extracellular Cu chelation with BCS increased the expression only of *Tc*IT by 2-fold but no effect on the expression of the other three genes was observed. These results can be summarized as *Tc*FR and *Tc*Fet3 genes did not respond to changes in Cu concentration in the epimastigote stage. *Tc*CuATPase slightly responded to changes in Cu availability, presenting significant differences when comparing extra Cu (lower mRNA level) and BCS (higher mRNA level) conditions. The most appreciable response was observed for *Tc*IT gene, whose mRNA levels decreased when Cu concentration was elevated and *vice versa*, enlarged when Cu chelators were present (BCS).

**Figure 6.**
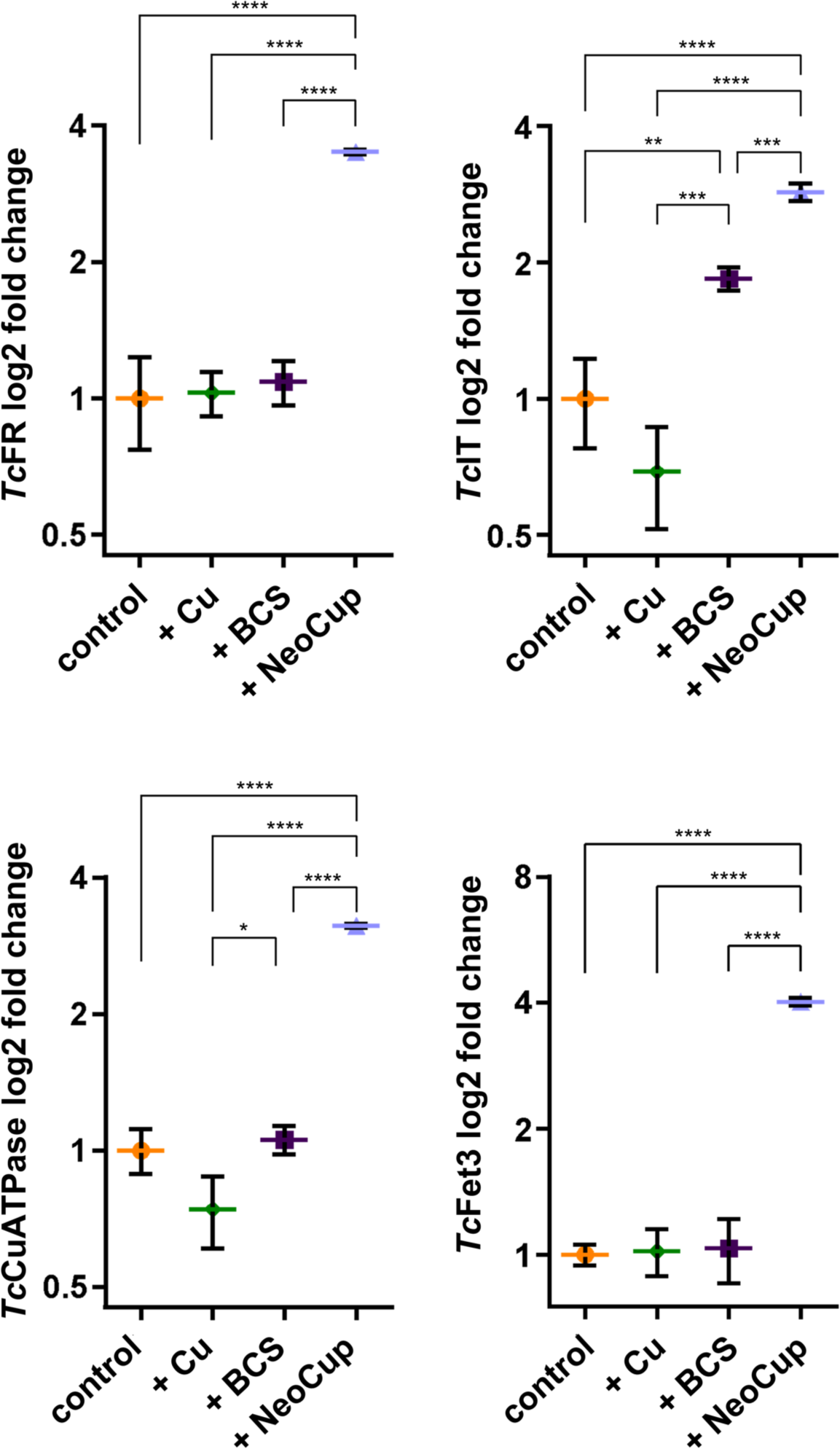
*Tc*IT and *Tc*CuATPase expression responses to Cu concentration in epimastigotes. Relative mRNA levels of *Tc*FR, *Tc*IT, *Tc*CuATPase, and *Tc*Fet3 genes in epimastigotes cultured for 2 days in LIT-10 % FBS-5 μM hemin medium supplemented with 250 μM copper (Cu), 500 μM bathocuproine disulfonate (BCS), and 1 μM neocuproine (NeoCup). The figure shows the Log2 of fold changes of the genes that were determined by 2^-ΔΔCt^ method. *Tc*Ubiquitin was used as housekeeping gene for normalization. The levels of mRNA of non-treated epimastigotes were used as reference (fold change = 1). The results are representative of at least two independent experiments in triplicate. Data represents averages ± SD. Differences were determined using one-way ANOVA and Bonferroni’s multiple comparisons post-hoc test. Significant differences are shown as asterisks: *p <0.05, **p <0.01, ***p <0.001, and ****p <0.0001.

### Copper is required in the differentiation process from epimastigotes to metacyclic trypomastigotes (metacyclogenesis)

To test the impact of Cu during metacyclogenesis, epimastigotes were incubated with different concentrations of Cu and Cu chelators for 7 days. The percentage of metacyclic trypomastigotes was determined for the different conditions in late stationary phase cultures (Fig. 7). The addition of 100 μM Cu did not affect the percentage of metacyclic trypomastigotes compared to the control. However, the addition of Cu chelators – 100 μM BCS or 0.5 μM NeoCup – significantly reduced the percentage of metacyclic trypomastigotes obtained to 40 % and 62 % of the control, respectively. A higher concentration of Cu (250 μM) affected the percentage of trypomastigotes but also significantly affected the number of total cells, so we could not differentiate between an effect on epimastigote replication or on metacyclogenesis *per se*. These data suggest that Cu plays a relevant role during the differentiation process from epimastigotes to metacyclic trypomastigotes, where any imbalance in the intracellular Cu availability negatively affects the metacyclogenesis.

**Figure 7.**
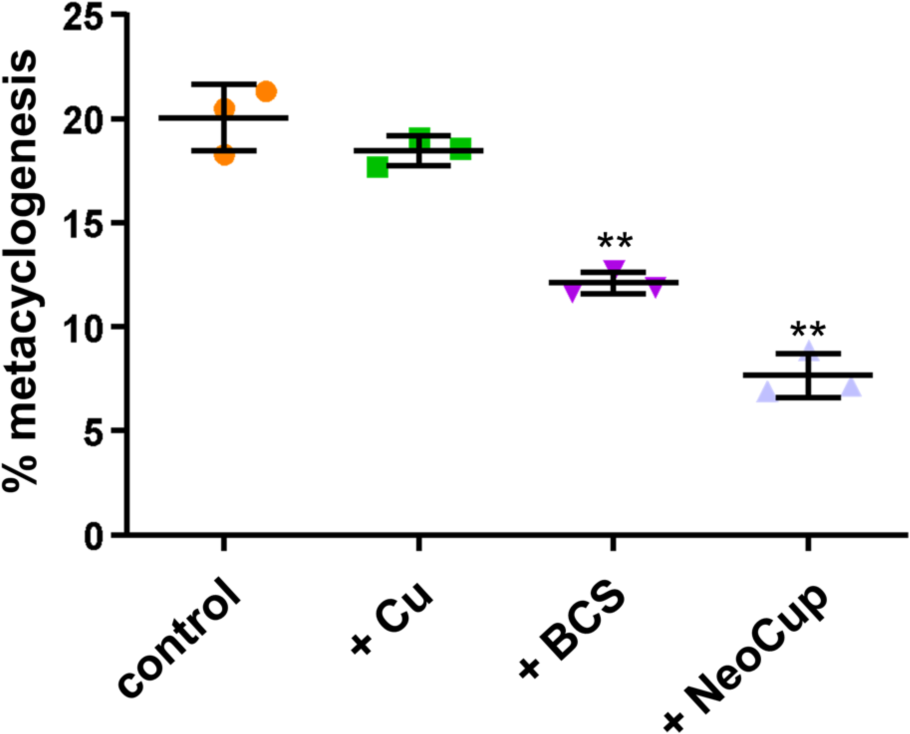
Copper chelators reduce metacyclogenesis differentiation. Percentage of metacyclic trypomastigotes in culture after 7 days incubation in LIT-10 % FBS-5 μM hemin pH 6 medium plus 100 μM copper (Cu), 100 μM bathocuproine disulfonate (BCS), 0.5 μM neocuproine (NeoCup), or no addition (control). The results are representative of at least two independent experiments in triplicate. Data represents averages ± SD. Differences compared with control were determined using one-way ANOVA and Dunnett post-hoc test. Significance is shown as asterisks: **p <0.001.

### Amastigotes experience copper stress during the intracellular replication

The effect of Cu was tested during the *T. cruzi* infection of Vero cells. Before the infection assay Vero cells were tested for maximal Cu tolerance during 72 h. Cell cultures did not show substantial changes when exposed to up to 200 μM Cu (tested range from 100 to 500 µM) and at least 500 µM BCS, but NeoCup was significantly cytotoxic at concentrations of 0.1 and 0.5 μM (Fig. 8). Then, parasites were incubated with Cu (50 μM, 100 μM, and 150 μM), BCS (500 μM), or NeoCup (0.005 μM, 0.01 μM, and 0.02 μM) during the whole experiment which design included the infection process with cell-derived trypomastigotes, their intracellular differentiation, and amastigote replication inside Vero cells. The percentage of infected cells (Fig. 9A-C) and the number of amastigotes per infected cell (Fig. 9D-F) were determined. Neither Cu nor Cu chelators showed a significant effect on infection (Fig. 9A-C). On the contrary, amastigote replication was affected by Cu and NeoCup but not BCS (Fig. 9E). Cu significantly reduced the number of amastigotes per infected cell (Fig. 8D) and NeoCup significantly increased it (Fig. 9F). Based on these results, we decided to analyze the involvement of *Tc*FR, *Tc*IT, *Tc*CuATPase, and *Tc*Fet3 genes during the response to Cu in amastigote stage by measuring changes in mRNAs levels resulting from Cu and NeoCup treatments (Fig. 10). *Tc*FR expression did not change with any treatment, but the other three genes increased their expression 2 times under Cu addition, and only *Tc*IT and *Tc*CuATPase were affected by the NeoCup treatment (1.5- and a 2-fold increases, respectively).

**Figure 8.**
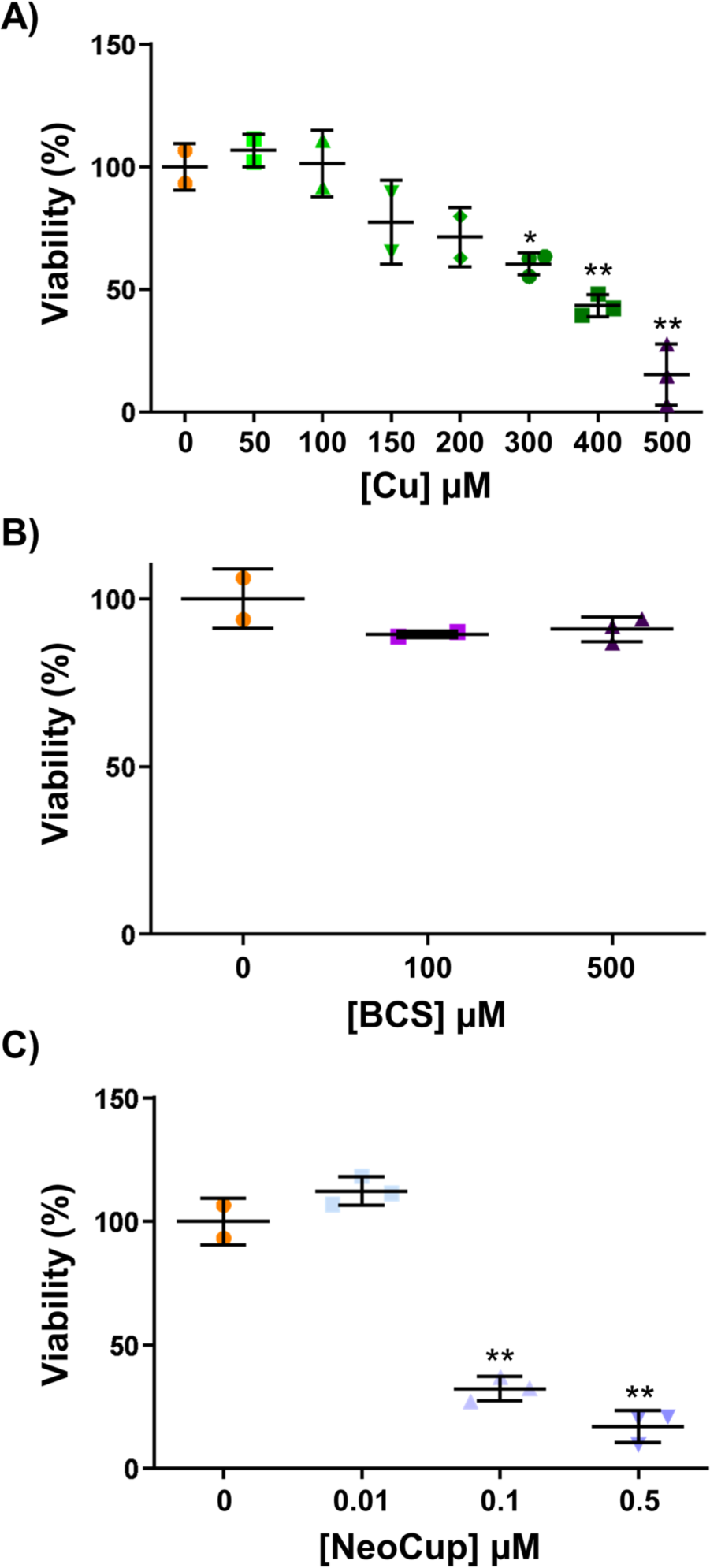
Vero cells tolerate high concentration of copper and BCS. Vero cells were incubated for 72hours in DMEM medium at 37°C with different concentrations of copper (Cu) (A), different concentrations of the chelator bathocuproine disulfonate (BCS) (B), and different concentrations of the chelator neocuproine (NeoCup) (C). After the incubation, the cytotoxicity was measured with MTT viability assay. The results are representative of at least two independent experiments in triplicate. Data represents averages ± SD. Differences were identified using one-way ANOVA and Dunnett post-hoc test compare with control. Significance is shown as stars: *p <0.01 and **p <0.001.

**Figure 9.**
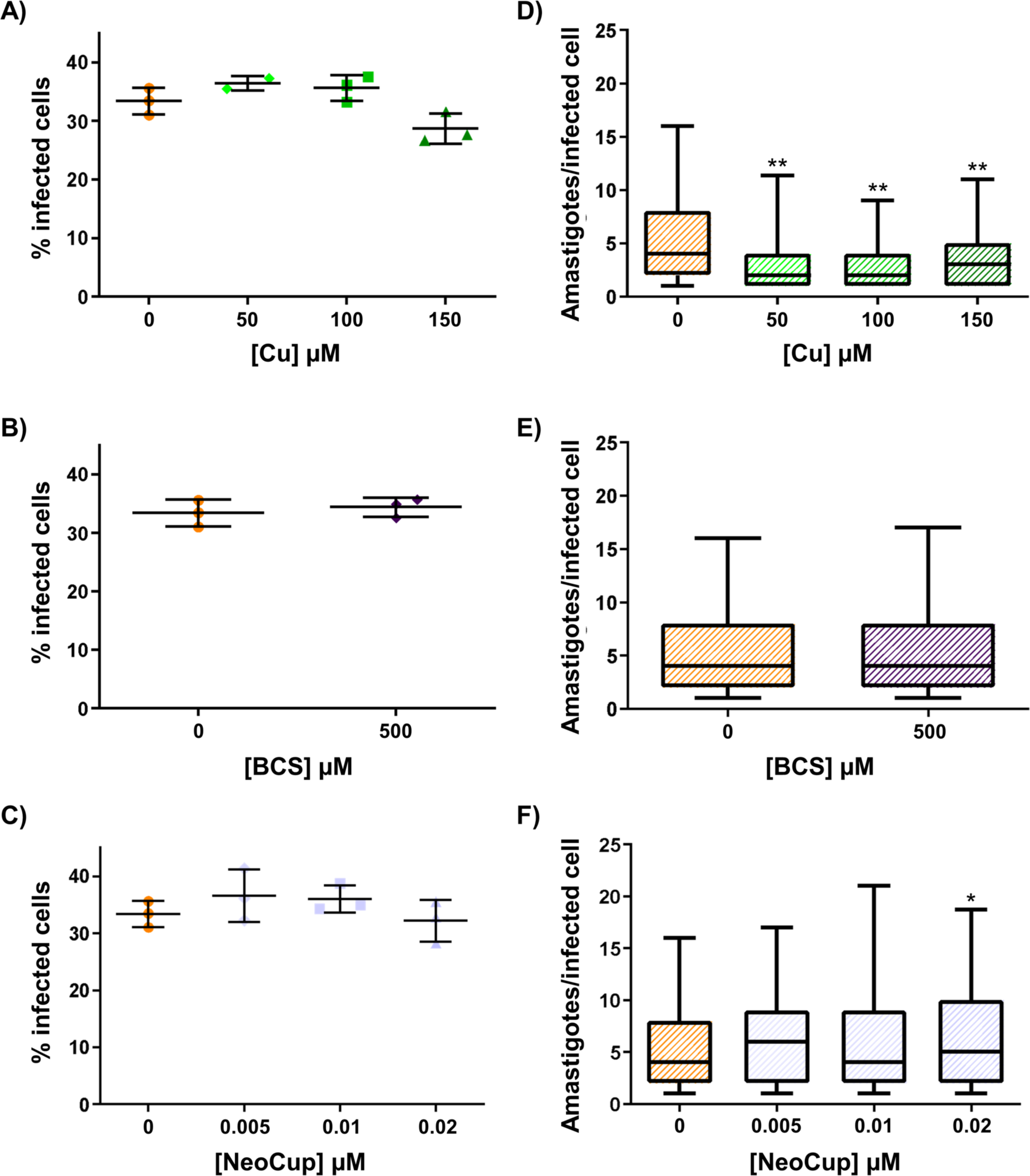
Copper affects intracellular amastigotes replication. Vero cells plated on cover glasses were incubated with cell-derived trypomastigotes (MOI of 10) for 16 hours in DMEM medium at 37°C plus different concentrations of copper (Cu) (A and D), the chelator bathocuproine disulfonate (BCS) (B and E), and different concentrations of the chelator neocuproine (NeoCup) (C and F). After that, the monolayered Vero cells were washed twice with PBS and the same medium was added but without trypomastigotes. After 72hours, the percentage of infected cells and the number of amastigotes per infected cell was evaluated. The results are representative of at least two independent experiments in triplicate. A-C) The percentage of infected cells is expressed as averages ± SD and no difference was identified using one-way ANOVA and Dunnett post-hoc test compares with control. (D-F) The number of intracellular amastigotes per infected cell is shown in a box and whisker plot (the box indicated the first quartile, the medium and the third quartile, and the whisker indicated the 5th and 95th percentile), and differences were identified using Kruskal–Wallis test and Dunn’s multiple comparison post-hoc test compare with control. Significance is shown as stars: *p <0.05 and **p <0.001.

**Figure 10.**
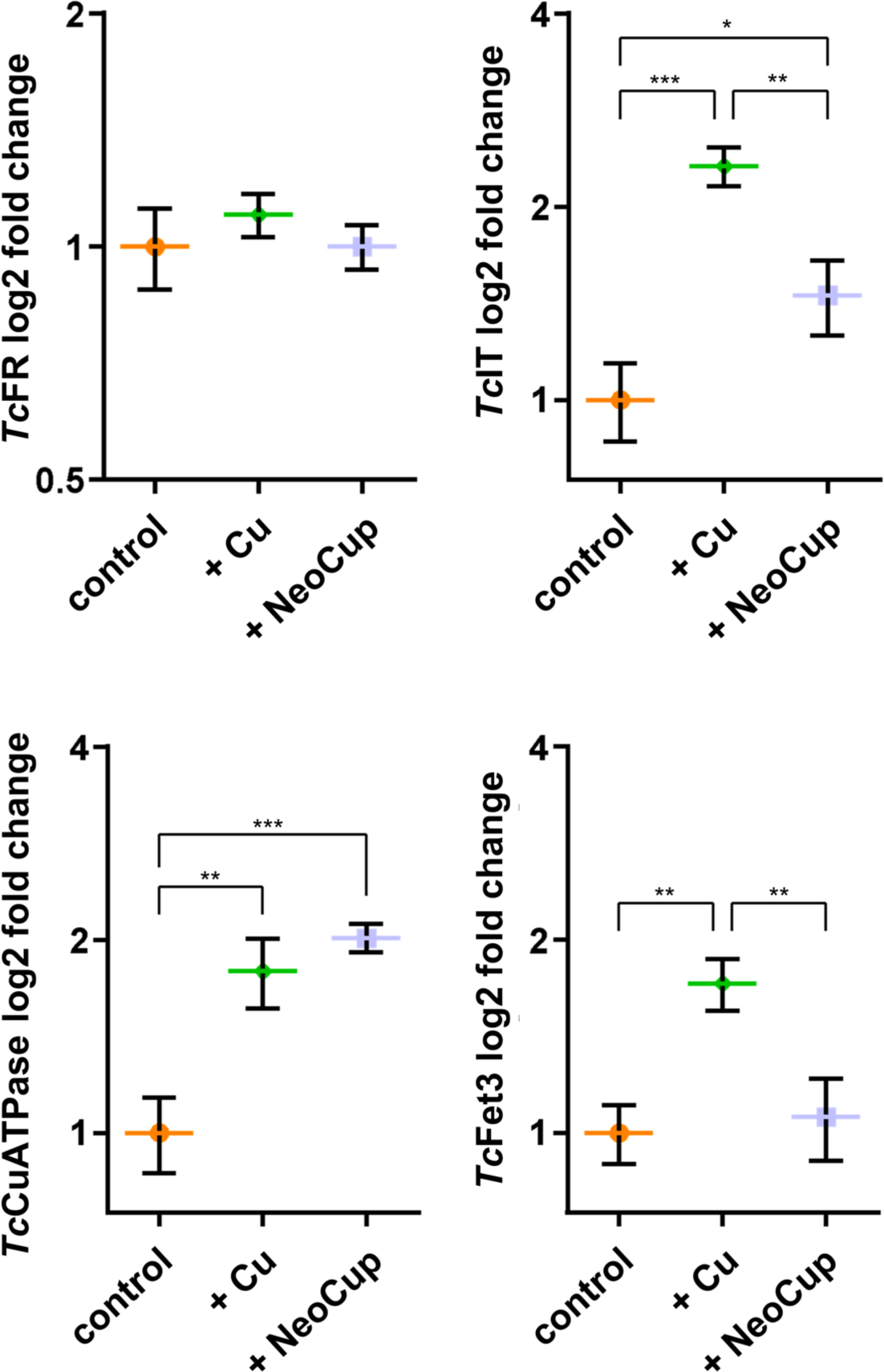
*Tc*IT, *Tc*CuATPase, and *Tc*Fet3 expression responses to Cu concentration in amastigotes. mRNA level of *Tc*FR, *Tc*IT, *Tc*CuATPase, and *Tc*Fet3 genes in amastigotes obtained from infected Vero cells maintained for 3 days in DMEM-2 % FBS supplemented with 100 μM copper (Cu) or 0.02 μM neocuproine (NeoCup). The figure shows the log2 of fold change of the genes that were determined by 2^-ΔΔCt^ method. *Tc*Ubiquitin was used as housekeeping gene for normalization. The no treated amastigote stage (fold change = 1) was used. The results are representative of at least two independent experiments in triplicate. Data represents averages ± SD. Differences were identified using one-way ANOVA and Bonferroni’s multiple comparisons post-hoc test. Significance is shown as stars: *p <0.05, **p <0.01, and ***p <0.001.

## Discussion

Copper is an essential cofactor for aerobic metabolism and at the same time it is a toxic ion for the cell. Copper trafficking, storage and utilization must be fine tune regulated and it has not been studied in *Trypanosoma cruzi.* This report is the first approach to understand *T. cruzi* copper metabolism and we demonstrated copper double-edged role: it is necessary for some processes such as growth and differentiation of epimastigotes, but it stresses intracellular amastigotes. We also identified the genes that are possibly involved in *T. cruzi* Cu homeostasis, which are summarized in Table 1, supporting our hypothesis by measuring these gene expression levels under changes in Cu concentration.

When testing the effect of Cu stress on the replicative stages (epimastigote and amastigote) as well as during metacyclogenesis, we found that epimastigotes tolerated high Cu concentrations and that Cu also was relevant for cellular function and metacyclic differentiation. However, intracellular amastigotes succumbed to Cu stress more easily than the other stages, the non-intracellular forms. Epimastigote tolerance and control of intracellular Cu content indicates that *T. cruzi*, at least at this stage, can regulate Cu transport to cope with high Cu exposure and probably to deliver this ion to its target cupro-proteins. Both processes must be mediated by dedicated proteins.

Regarding Cu restriction by the addition of Cu chelator to epimastigote cultures, only high concentrations of BCS, that cause a drop in intracellular Cu concertation, moderately affected the growth, without any effect on oxygen consumption. Whereas the NeoCup treatment markedly decreased the epimastigote growth together with only a 20 % decrease in oxygen consumption. We hypothesize that NeoCup is affecting also intracellular Cu bioavailability and that the drop of attainable Cu affected essential cellular processes in the parasite with little or no effect in mitochondrial functions or that under Cu withdrawal, the parasite prioritizes essential mitochondrial respiration over other metabolic functions. These observations suggest that Cu is also involved in essential processes for epimastigote growth other than mitochondrial respiration. The fact that epimastigotes were able to tolerate high and low concentrations of Cu, reinforce the hypothesis that *T. cruzi* is able to adapt its metabolism to changes in the availability of this cofactor maintaining its intracellular concentration in a sustainable range.

It is worth noting that *T. cruzi* replicative stage in its mammalian host (i.e., amastigote) is an intracellular form of the parasite. Circulating infective flagellated trypomastigotes are internalized by the mammalian host cells where they should escape the parasitophorous vacuole to the cytosol (of macrophages, fibroblast, epithelial cells, as a first step of parasite invasion) to be able to transform into amastigotes [5,27]. During this process, *T. cruzi* must face the innate immune host response. Cu has been shown to have a fundamental role in defense as described also for other intracellular pathogens for example *Mycobacterium tuberculosis*, or fungus, such as *Cryptoccocus neoformans* [28,29]. In this sense, our data show that Cu supplementation during Vero cell infection *in vitro* reduced amastigote proliferation while the presence of Cu chelators augmented it. These results support the hypothesis that *T. cruzi* must evade the impact of Cu as part of the host defense mechanisms against the parasite during the infection process.

In eukaryotic organisms, control of Cu homeostasis involves proteins at different localizations: plasma membrane, cytosol, and organelles. In the plasma membrane, there are copper reductases and copper transmembrane transporters. In the cytosol, there are copper chaperones that transport Cu from the plasma membrane to cytosolic cupro-proteins, such as superoxide dismutase (SOD), or to organelles. The main target organelles are *trans*-Golgi complex and mitochondria [1]. Many of these proteins have been identified as conserved in a wide range of eukaryotes but the survey of the trypanosomatids is less complete. We identified several of these proteins encoded in the *T. cruzi* genomic sequence, which are listed in Table 1. Our findings suggest that *T. cruzi* has conserved the mitochondrial proteins such as *Tc*Sco1, *Tc*Sco2, *Tc*Cox11, *Tc*Cox17, *Tc*Cox19, and the cytochrome c oxidase homolog (recently revised in [30]) but it also presents some potentially novel members such as *Tc*Sco17-like and *Tc*Sco-like. In addition, a conserved P-type ATPase (*Tc*CuATPase) likely responsible for transport into the Golgi network for delivery to targets and probably responsible for the tolerance to Cu stress was identified. Our bioinformatic search in *T. cruzi* genome (TritrypDB, http://tritrypdb.org/tritrypdb/) identified 3 ORFs coding for CuATPases in the Dm28c strain genome. The three genes encode for 954 amino acid proteins and present more than 99.6 % identity between them. The last genome sequence reported [31] was determined through third generation long-read sequencing technologies allowing a more precise ascertainment of the number of genes present confirming that several copies are indeed present (C4B63_19g251, C4B63_19g252, C4B63_19g254, and C4B63_19g256). The fact that the three (or four) *Tc*CuATPase gene copies are arranged in tandem and do not constitute an error during the sequence assembly process could suggest functional role for different variants of the protein or that these genes have yet to decay over time. In contrast, no homologs of the canonical plasma membrane transporters were identified in our searches. Besides the mitochondrial proteins and the *Tc*CuATPase, we identified other genes encoding for proteins whose function would be related with control of Cu homeostasis, *Tc*IT, *Tc*FR, and *Tc*Fet3 (Table 1).

We analyzed if this proposed transport machinery composed of *Tc*CuATPase, *Tc*IT, *Tc*FR, and *Tc*Fet3 were transcribed in different life cycle stages. The four genes were differentially expressed in all life cycle stages studied. The results suggested that they are relevant in the infective stages although the expression patter is not identical for all of them. Different expression of *Tc*IT between metacyclic trypomastigote and epimastigote stages were reported previously when *Tc*IT was studied as Fe transporter by Dick and colleagues. Also, the expression of these genes under Cu stress were analyzed and they showed different behaviors. The expression of *Tc*FR gene did not respond to any treatment in the replicative stages (epimastigotes and amastigote), except for NeoCup in epimastigotes. *Tc*Fet3 expression is up-regulated only after Cu addition in amastigote stage and NeoCup addition in epimastigotes. Instead, *Tc*IT and *Tc*CuATPase responded to increment in intracellular Cu in the replicative stages, but the response was opposite between the two stages being up- regulated in amastigotes and down-regulated in epimastigotes. Both genes increased their expression under the presence of Cu chelators (BCS and NeoCup) in epimastigotes and NeoCup in amastigotes. The different response of these two genes is consistent, at least in part, with the different phenotypes observed in the epimastigote growth and the amastigote replication under stress by excess or defect of Cu. Despite this, we postulate that *Tc*IT and *Tc*CuATPase may be involved in Cu response to modulate the intracellular Cu concentration. Lastly, the differences observed between BCS and NeoCup could be related to the fact that the last one is able to freely traverse plasma membranes [32] so it can sequester intracellular Cu in comparison to BCS that would lower culture medium Cu concentrations. NeoCup produced a stronger effect in Cu availability, in cellular homeostasis and in transcript levels.

It is well stablished that a link exists between Cu and Fe transport because they share homeostasis machinery and because multi-copper oxidases are critical to Fe transport [14,16]. Considering that Cu and Fe could share transporters (as divalent cation transporters), we hypothesized that Cu could affect Fe uptake or *vice versa* in *T. cruzi*. While Cu overload decreased Fe content in epimastigotes, it did not affect heme content, reinforcing the idea of a shared transporter of the divalent cations. The addition of Fe and heme to the medium did reduce the Cu content of parasites further supporting an interplay or competition between these ions. This is consistent with *Tc*FR and *Tc*IT regulation proposed by Dick and colleagues [33], where they showed that the transcript levels of *Tc*FR and *Tc*IT responded to modulation of Fe and heme content of the media in cultures of *T. cruzi* epimastigotes: *Tc*FR was upregulated when heme content was depleted but did not respond to Fe concentration, and *Tc*IT was downregulated at high heme and Fe concentration. According to our data, Cu chelators did not influence Fe content, even though they caused *Tc*IT upregulation. This could be explained if other genes involved in Fe uptake (or heme as source of Fe), as *Tc*FR, are already expressed and do not increase their expression with chelator treatments. This is congruent with the fact that heme cause a higher decrease on intracellular Cu than Fe and could be explained if Cu transport depends on *Tc*FR instead *Tc*IT. On the other side, Cu overload reduced intracellular Fe but not heme consistent with our data that the presence of this ion reduced the expression of *Tc*IT but not *Tc*FR. Despite that we postulate the importance of *Tc*IT and *Tc*CuATPase, other genes could be involved in the Cu and Fe uptake and further studies are necessary to elucidate the regulation mechanism that explain the observed contrarieties.

Interestingly, Cu addition allowed a growth enhancement of heme starved epimastigotes and elevated their superoxide levels. We relate both observations because the heme was reported to activate the proliferation of parasites mediated by ROS production and the activation of calcium-calmodulin-dependent kinase II-like cascade [34] and, as well, Cu is known to produce ROS bursts in other organisms [1]. Then, this transient growth rescue could be due to ROS production by Cu while parasites can use stored heme. About metacyclogenesis, it was negatively affected by a Cu deprivation that is consistent with results obtained by Dick and colleagues, where they observed that Fe depletion reduced this process [33]. In agreement with that, ROS (hydrogen peroxide) is also involved in *T. cruzi* amastigogenesis (differentiation from trypomastigotes to amastigotes) [35] . Our hypothesis is that Cu could be involved in the differentiation processes because it is an essential cofactor for it, or Cu produces a necessary oxidative signal in the parasite.

The finding that expression levels of the four genes studied were lower in replicative stages than in infective ones could be interpreted as a way of modulation of Cu uptake and homeostasis depending on the levels of this ion encountered in the environments the parasite faces in each life cycle stage. In other words, epimastigotes in the triatomine gut and amastigotes inside the mammalian host cell must support higher local concentrations of Cu and, a way to control the entrance of it, could be by reducing the expression of its transporters, chaperones and other proteins involved in this ion management. This type of regulation is observed for *Tc*HRG [36,37] whose expression is modulated according to the heme availability in the growth medium until the intracellular *quota* of this cofactor is achieved. Reinforcing this idea, when epimastigotes were *in vitro* confronted to Cu stress, we found that *Tc*FR, *Tc*Fet3, *Tc*IT and *Tc*CuATPase increased with NeoCup and that *Tc*IT and *Tc*CuATPase also responded to BCS and Cu. Based on the data presented here, we proposed a unique network for Cu uptake at the plasma membrane, probably at the flagellar pocket region, and distribution in the cytosol, which is schematized in Fig. 11. In this model, *Tc*FR reduces Cu^2+^ to Cu^+^ and *Tc*IT transports it across the plasma membrane. The molecule trypanothione (TrySH) intracellularly distributes and stores Cu. *Tc*CuATPase transports Cu in the endomembrane system, probably to the multicopper oxidase *Tc*Fet3, and/or exports copper outside the cell. In the mitochondrion, the transmembrane transporter *Tc*Pic2 and the chaperones *Tc*Sco1, *Tc*Sco2, *Tc*Sco-like, *Tc*Cox11, *Tc*Cox17, *Tc*Cox17-like, and *Tc*Cox19 uptake and deliver copper to the cytochrome *c* oxidase (COX) of the electron transport chain.

**Figure 11.**
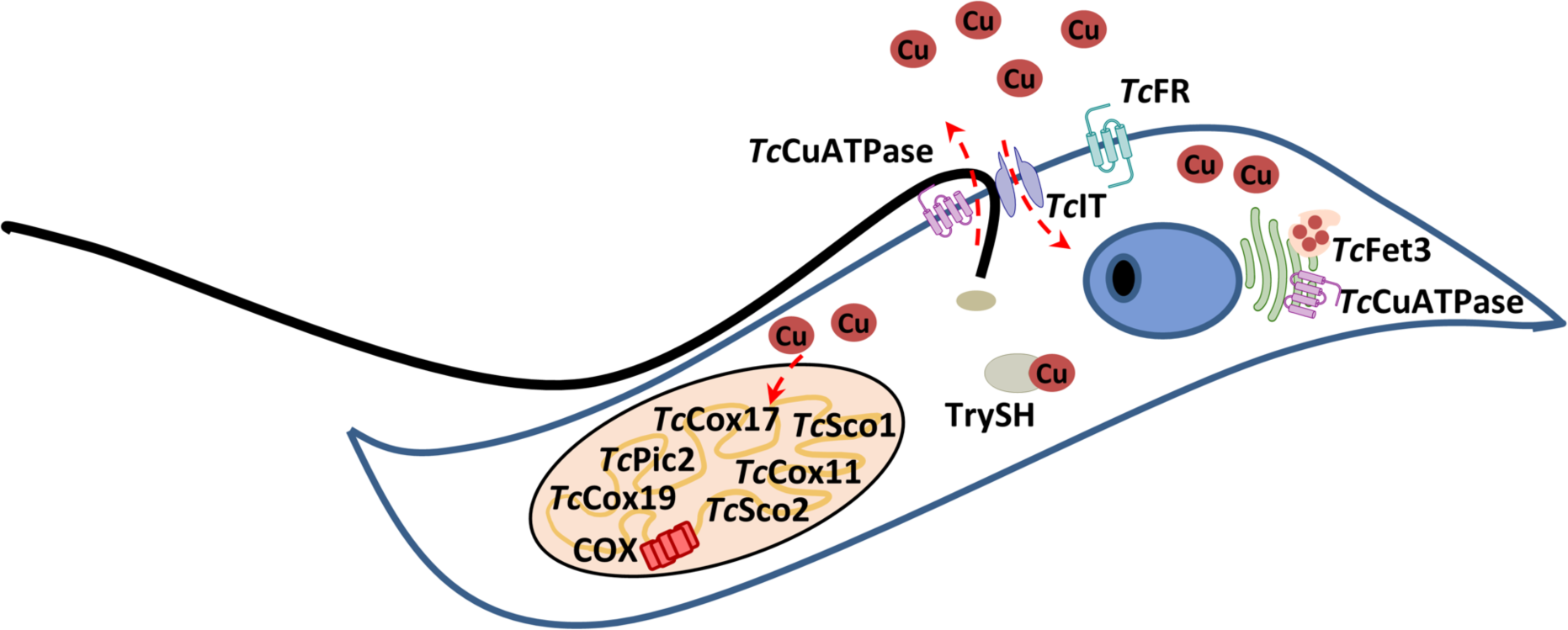
Proposed model of copper uptake and distribution in *T. cruzi*. Copper (Cu) could be imported by the transmembrane iron transporter *Tc*IT after its reduction by the ferric reductase *Tc*FR. Trypanothione (TrySH), analogue to glutathione in trypanosomatids, could be involved in intracellular copper trafficking and storage. The transmembrane transporter *Tc*CuATPase could be involved in copper transport in endomembrane system and/or exportation of it. The multicopper oxidase *Tc*Fet3 could receive Cu from the *Tc*CuATPase. In the mitochondrion, several proteins could participate in copper uptake and delivery to the cytochrome *c* oxidase (COX), as the transmembrane transporter *Tc*Pic2 and the chaperones *Tc*Sco1, *Tc*Sco2, *Tc*Sco-like, *Tc*Cox11, *Tc*Cox17, *Tc*Cox17-like, and *Tc*Cox19. No cytosolic chaperones proteins were identified, as homologues to the yeast copper chaperones Atx1 and Ccs; neither high affinity membrane copper transporters as homologues to the yeast Ctr1, Ctr2, or Ctr3; or homologues to the copper/zinc superoxide dismutase SOD1.

Overall, this report documents the roles for copper in the life cycle of the parasite, identifies potential copper homeostasis proteins (Table 1 and Fig. 11), and provides the evidence of gene regulation. Further studies are necessary to understand the full set of phenotypes exhibited during Cu stress, such as the determination of the exact localization of these proteins in *T. cruzi* or finding the missing factors as the cytosolic chaperones. As this report demonstrates, copper metabolism must be researched in *T. cruzi* because it is important along its life cycle, especially during the infection. Understanding key metabolism is essential to develop new strategies against this pathogen.

## Materials and Methods

### Reagents

Dulbecco’s Modified Eagle Medium (DMEM) was obtained from Life Technologies Corporation (Grand Island, NY, USA). Fetal Bovine Serum (FBS) was obtained from Internegocios S.A. (Buenos Aires, Argentina) and heat-inactivated at 56 °C for 30 min. Hemin stock solution (1 mM) was prepared in 50 % (v/v) EtOH, 0.01 N NaOH, fractionated and stored at −80 °C. Copper (Cu) was added as Cu^2+^ from CuSO4 salt. Bathocuproine disulfonate (BCS, an extracellular copper chelator) was obtained from Sigma-Aldrich (St. Louis, MO, USA). Neocuproine solution (NeoCup, an intracellular copper chelator) was prepared from Neocuproine hydrochloride monohydrate (Supelco, St. Louis, MO, USA). MTT (3-(4,5- dimethylthiazol-2-yl)-2,5-diphenyltetrazolium bromide) was obtained from Sigma-Aldrich, St. Louis, MO, USA)

### In silico analysis

The searches in the TriTrypDB (http://tritrypdb.org/tritrypdb/) (Amos *et al.*, 2022) were done using known amino acid sequences of Cu transporters, chaperones, and cupro-proteins in other organisms as seeds (Table 1). The identified sequences were analyzed also with Conserved Domain Database (https://www.ncbi.nlm.nih.gov/Structure/cdd/wrpsb.cgi) (Lu *et al.*, 2020) to verify conserved domains and all of them were confirmed (Table 2). The sequences from *Saccharomyces cerevisiae*, used as seeds, were obtained from *Saccharomyces* Genome Database (SGD, www.yeastgenome.org) [38]. The other sequence seeds were obtained from National Center for Biotechnology Information (NCBI) (https://www.ncbi.nlm.nih.gov/protein/). TrypTag database (http://tryptag.org) [39] was used for the subcellular localization data of the *T. brucei* orthologs. When no homologues were found in the TriTrypDB, the negative result was confirmed with psi-BLAST search from NCBI (https://blast.ncbi.nlm.nih.gov) and with FoldSeek [40] searches.

**Table 2.**
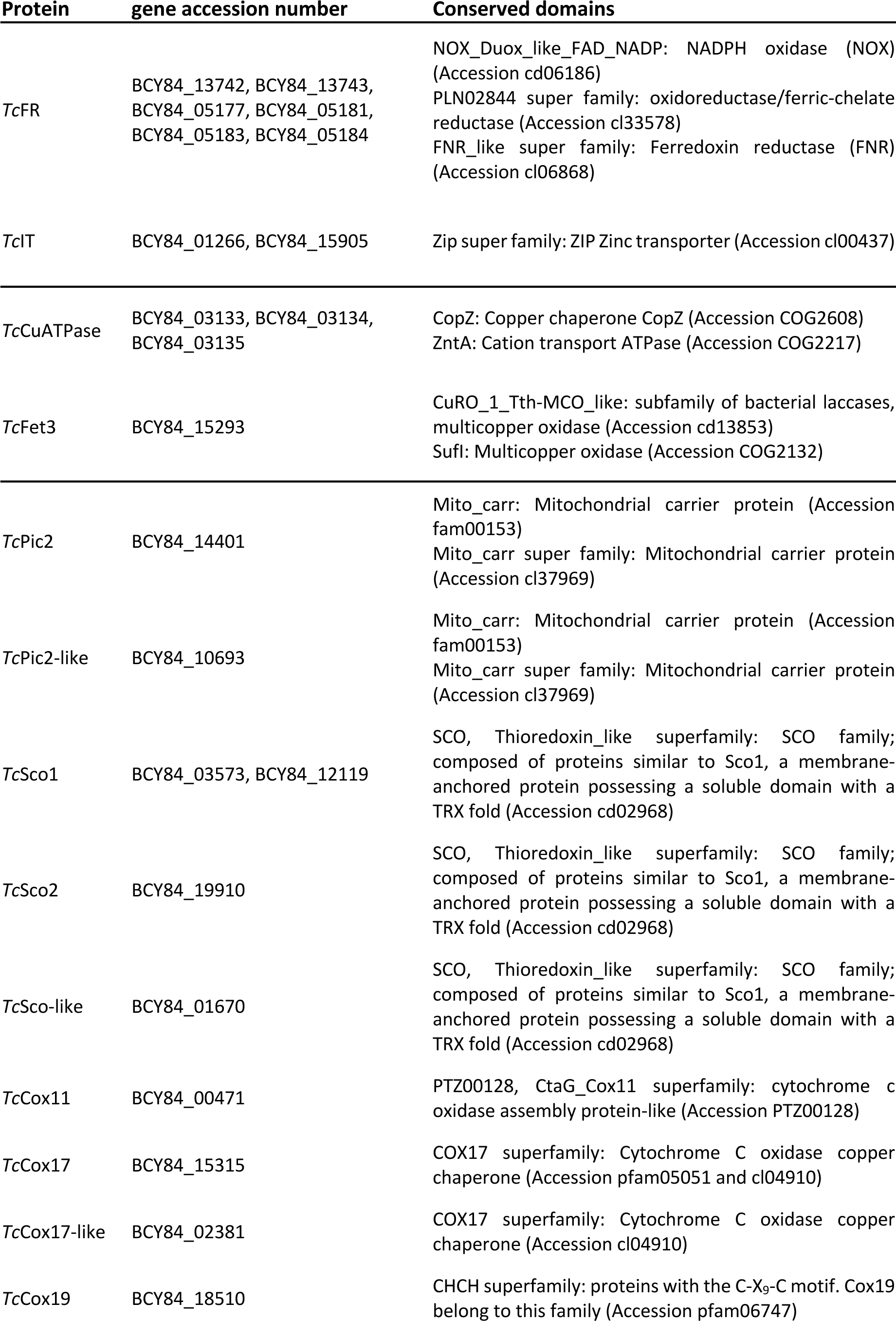
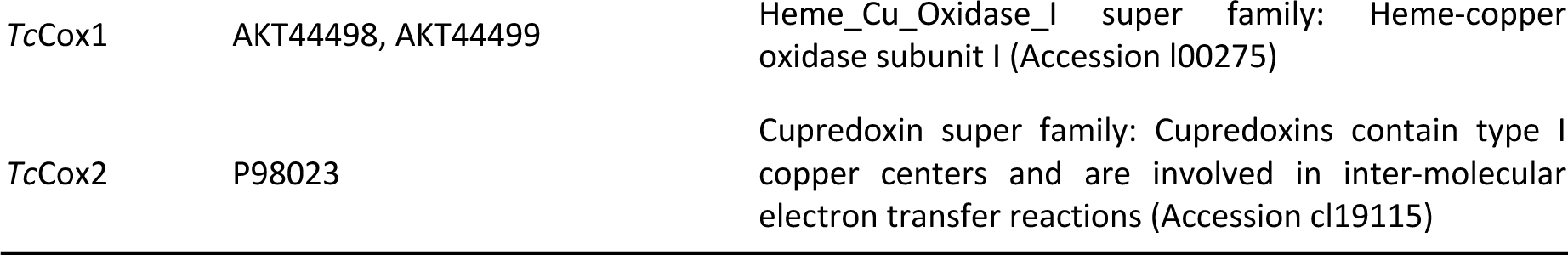
Conserved domains identified by Conserved Domain Database. Conserved Domains of the identified *T. cruzi* sequences of Table 1. Expect Value threshold: 0.01

| Protein | gene accession number | Conserved domains |
| --- | --- | --- |
| *TcFR* | BCY84_13742, BCY84_13743,<br>BCY84_05177, BCY84_05181,<br>BCY84_05183, BCY84_05184 | NOX_Duox_like_FAD_NADP: NADPH oxidase (NOX)<br>(Accession cd06186)<br>PLN02844 super family: oxidoreductase/ferric-chelate<br>reductase (Accession cl33578)<br>FNR_like super family: Ferredoxin reductase (FNR)<br>(Accession cl06868) |
| *TcIT* | BCY84_01266, BCY84_15905 | Zip super family: ZIP Zinc transporter (Accession cl00437) |
| *TcCuATPase* | BCY84_03133, BCY84_03134,<br>BCY84_03135 | CopZ: Copper chaperone CopZ (Accession COG2608)<br>ZntA: Cation transport ATPase (Accession COG2217) |
| *TcFet3* | BCY84_15293 | CuRO_1_Tth-MCO_like: subfamily of bacterial laccases,<br>multicopper oxidase (Accession cd13853)<br>SufI: Multicopper oxidase (Accession COG2132) |
| *TcPic2* | BCY84_14401 | Mito_carr: Mitochondrial carrier protein (Accession<br>fam00153)<br>Mito_carr super family: Mitochondrial carrier protein<br>(Accession cl37969) |
| *TcPic2-like* | BCY84_10693 | Mito_carr: Mitochondrial carrier protein (Accession<br>fam00153)<br>Mito_carr super family: Mitochondrial carrier protein<br>(Accession cl37969) |
| *TcSco1* | BCY84_03573, BCY84_12119 | SCO, Thioredoxin_like superfamily: SCO family;<br>composed of proteins similar to Sco1, a membrane-<br>anchored protein possessing a soluble domain with a<br>TRX fold (Accession cd02968) |
| *TcSco2* | BCY84_19910 | SCO, Thioredoxin_like superfamily: SCO family;<br>composed of proteins similar to Sco1, a membrane-<br>anchored protein possessing a soluble domain with a<br>TRX fold (Accession cd02968) |
| *TcSco-like* | BCY84_01670 | SCO, Thioredoxin_like superfamily: SCO family;<br>composed of proteins similar to Sco1, a membrane-<br>anchored protein possessing a soluble domain with a<br>TRX fold (Accession cd02968) |
| *TcCox11* | BCY84_00471 | PTZ00128, CtaG_Cox11 superfamily: cytochrome c<br>oxidase assembly protein-like (Accession PTZ00128) |
| *TcCox17* | BCY84_15315 | COX17 superfamily: Cytochrome C oxidase copper<br>chaperone (Accession pfam05051 and cl04910) |
| *TcCox17-like* | BCY84_02381 | COX17 superfamily: Cytochrome C oxidase copper<br>chaperone (Accession cl04910) |
| *TcCox19* | BCY84_18510 | CHCH superfamily: proteins with the C-X<sub>9</sub>-C motif. Cox19<br>belong to this family (Accession pfam06747) |
| *TcCox1* | AKT44498, AKT44499 | Heme_Cu_Oxidase_I super family: Heme-copper oxidase subunit I (Accession I00275) |
| *TcCox2* | P98023 | Cupredoxin super family: Cupredoxins contain type I copper centers and are involved in inter-molecular electron transfer reactions (Accession cl19115) |

### RNA isolation and quantitative real-time PCR (qRT-PCR)

The RNA samples of the different life cycle stages of *T. cruzi* were obtained from: A) Epimastigotes cultured in LIT-10 % FBS-5 μM medium supplemented with 250 μM Cu, 500 μM BCS, or 1 μM NeoCup. After 2 days, the cells were collected. B) Metacyclic trypomastigotes were obtained as described for metacyclogenesis quantification. After 7 days, cells were collected and trypomastigotes were separated from epimastigotes using DEAE-cellulose as described (Cruz-Saavedra *et al.*, 2017). C) Cell-derived trypomastigotes were collected from the supernatant of infected Vero cells 5 days post infection. D) Amastigotes were obtained from infected monolayers of Vero cells maintained in DMEM-2 % FBS supplemented with 100 μM Cu or 0.02 μM NeoCup. At 72 h post infection, amastigotes were isolated as describe previously [7]. In each case, 100 x 10^6^ cells were used to RNA extraction using 1 mL TRI Reagent® (TR118 - Molecular Research Center, Inc., Cincinnati, OH, USA). The quality of RNA was verified in agarose gels and the RNA was quantified by absorbance. The RNA samples were treated with DNAse (RQ1 Rnasa-Free Dnase, Promega, Madison, WS, USA) and retrotranscribed with FireScript KIT Reverse Transcriptase (Solis BioDyne, Tartu, Estonia) to obtained cDNA. The quantitative Real Time PCR reactions were performed with HOT FIREPol EvaGreen qPCR Mix Plus (ROX) (Solis BioDyne, Tartu, Estonia), using the primers listed in Table S1 (Macrogen, Korea), in the CFX Opus 96 Real-Time PCR System (#12011319 - Bio-Rad, Miami, FL, USA) following the next protocol: 95 °C (10 min); 40 cycles of 95 °C (10 s), 60 °C (30 s), and 72 °C (20 s); followed by the denaturation curve to analyze the Tm of the products. The efficiency of each pair of primers was determined and the relative fold changes were determined by the 2^-ΔΔCt^ method [41]. The gene *Tc*Ubiquitin was used as reference (housekeeping) and the non-treated epimastigote stage was taken as reference (mRNA level = 1); except in the case of expression analysis of amastigote treatments where the non-treated amastigotes were used as reference.

### Parasites and cell lines

The experiments were done using *T. cruzi* Dm28c strain and the infections were performed in Vero cell line (ATCC CCL-81, available in the laboratory). *T. cruzi* epimastigotes were maintained in mid-log phase by periodic dilution in Liver Infusion Tryptose (LIT) medium (5 g/L liver infusion, 5 g/L bacto-tryptose, 68 mM NaCl, 5.3 mM KCl, 22 mM Na2HPO4, and 0.8 % (w/v) glucose, pH 7.4) supplemented with 10 % FBS and 5 μM hemin (LIT-10 % FBS-5 μM hemin medium), at 28 °C. Vero cells were cultured in DMEM plus 0.15 % (w/v) NaHCO_3_ and 10 % FBS (DMEM-10% FBS) at 37 °C in a humid atmosphere containing 5 % CO_2_.

### Epimastigote growth curves

Epimastigotes were routinely maintained at mid-log phase of growth by periodic dilution every 3 days in fresh LIT (LIT-10 % FBS-5 μM) medium for at least 10 days. Then, the cells were diluted at a concentration of 5 x 10^6^ parasites/mL and challenged to grow in LIT medium supplemented with 50-500 μM Cu, 50-500 μM BCS, or 0.01-100 μM NeoCup. The number of cells was monitored for 5-10 days. In the case of curves in LIT-10 % FBS-0 μM hemin medium, epimastigotes were cultured in this medium for at least 10 days before the treatments. The number of cells was monitored for 5-10 days. Cell growth was determined by cell counting in an automatized hemocytometer adapted to count epimastigotes (WL 19 Counter AA, Weiner Laboratory SAIC, Rosario, Santa Fe, Argentina) and in Neubauer counting chamber.

### Oxygen consumption measurements in epimastigotes

The oxygen consumption rates of epimastigotes (O_2_ nmoles/mL min 10^6^ cells) were quantified from the linear response, using a Clark electrode connected to a 5300 Biological Oxygen Monitor (Yellow Springs Instrument Co.), as previously described [7,42].

### Superoxide anion production in epimastigotes

Superoxide anion production was quantified as it was previously described in [43] with some modifications. Briefly, epimastigotes were maintained for 10 days in mid-log phase by periodic dilutions in LIT-10 % FBS without hemin medium. Then, the cells were diluted at a concentration of 5 x 10^6^ parasites /mL and challenged to grow in LIT-10 % FBS without hemin medium plus 100 µM Cu or no addition as control. After 5 days, 50 x 10^6^ cells per sample were collected, washed with PBS, resuspended in 270 µL PBS plus 30 µL of 10 mg/mL NBT (nitro- blue tetrazolium chloride), and incubated 30 min at 28 °C. After the incubation, the cells were washed with PBS and lysed with 161.6 µL DMSO. The mix was vortexed and then 138.4 µL of 2 M NaOH were added. Absorbances of the samples were measured in microplate reader (Multimodal Microplate Reader, model Synergy HTX version S1LFA, BioTek Instruments, Shoreline, WA, USA) at 540 nm and the concentration of superoxide anion was calculated using an extinction coefficient of 7.2 cm^2^ µmol^−1^.

### Cu and Fe contents determined by ICP-OES

Cu and Fe contents were determined by ICP-OES as described in [24]. Briefly, epimastigotes were maintained for 6 days in LIT-10 % FBS-5 μM hemin medium plus different concentrations of copper or copper chelators, or in LIT-10 % FBS without hemin medium plus different concentrations of hemin or iron. For heme starved cultures, the parasites were maintained the 10 previous days at mid-log phase by periodic dilutions in fresh LIT-10 % FBS without hemin medium before the treatments. 120 x 10^6^ epimastigotes were collected per measurement and lyophilized. The samples were digested by boiling for 1 h in 40 % nitric acid, then diluted in ultra-pure metal-free water and analyzed by ICP-OES (Perkin Elmer, Optima 7300DV) *versus* acid-washed blanks. Concentrations were determined from a standard curve constructed with serial dilutions of two commercially available mixed metal standards (Optima). Blanks of nitric acid with and without ‘metal-spikes’ were analyzed to ensure reproducibility.

### Heme content determined by pyridine method

The heme content was measured by the alkaline pyridine method described in [44]. We made modifications to perform the assay with epimastigotes that were described in [45]. Heme concentration was calculated from the differential -reduced (by adding sodium dithionite) minus oxidized- spectrum. Molar extinction coefficient at 557 nm: 23.98 mM^−1^ cm^−1^.

### Quantification of metacyclogenesis

Metacyclic trypomastigotes were obtained from epimastigotes (late stationary phase) as previously reported [46]. The medium was supplemented with Cu, BCS or NeoCup. The cells were collected and fixed with 3.7 % (w/v) formaldehyde in PBS. Then, the number of epimastigotes and metacyclic trypomastigotes were counted in Neubauer chamber.

### Cytotoxicity assays on Vero cells

Vero cells were plated on 96 multi-well plate in DMEM-2 % FBS (4 x 10^3^ cells per well) and incubated for 24 hours. Then, the cells were washed with PBS and incubated with 200 μL of DMEM-2 % FBS supplemented with 50-500 μM Cu, 50-500 μM BCS, or 0.01-5 μM NeoCup. No addition was made in the 100 % viability control. After 72 h, the viability of cells was measured by MTT reduction colorimetric method as described [47]. Briefly, the cells were incubated in 180 μL DMEM-2 % FBS plus 20 μL 5 mg/mL MTT for 1 h, washed with PBS and mixed with 100 μL DMSO. Absorbance at 540 nm was measured to determine formazan production.

### Infection assay

Vero cells were plated on 24-well plates with coverslips, 3 x 10^3^ cell/well, and incubated in DMEM- 2 % FBS. After 24 h, they were incubated with cell-derived trypomastigotes (Multiplicity of Infection, MOI of 10) for 16 h in DMEM-2 % FBS supplemented with 50-150 μM Cu, 50-500 μM BCS, or 0.005-0.02 μM NeoCup. Cell-derived trypomastigotes were obtained previously from an infection of monolayered Vero cells with metacyclic trypomastigotes. Then, Vero cells were washed twice with PBS and the medium was renewed but without trypomastigotes. 72 h post infection, the monolayers of infected Vero cells were washed with PBS, fixed with methanol for 15 min, washed with stabilized water, and stained with Giemsa’s reagent for 30 min. Then, they were washed with stabilized water and mounted with Canada balsam. 300 Vero cells per slide were analyzed under the microscope to determine the percentage of infected cells and the number of amastigotes per infected cell.

### Statistical analysis

The assays were independently performed at least two times, with its biological and technical replicas. The expression of results and the appropriate statistical analysis used in each case is informed at the corresponding figure legend. All the analyses were done with Prism v 6.0 software (GraphPad Software, San Diego, CA, USA).

## Acknowledgments

We are grateful to laboratory technician Dolores Campos for technical assistance in cell culture procedures. The research leading to these results has received funding from the National Agency of Scientific Investigation, Technological Development, Promotion, and Innovation (grant number PICT 2018-02110 to JAC, from 2020 to 2023), Agencia Santafesina de Ciencia, Tecnología e Innovación (ASaCTeI) (grant number PEIC I+D 2021-130 to JAC, from 2022 to 2023). MLM was supported by a postdoctoral fellowship from the National Scientific and Technical Research Council (CONICET).

## Author contributions

JAC and MLM conceived and designed the project; MLM performed most of designed experiments; MGM performed qRT-PCR assays; XZ: performed ICP-OES experiments; MLM, MGM and JAC prepared figures; JAC, MLM, MGM and PAC discussed results. JAC supervised the project; JAC, MLM, MGM and PAC wrote and edited the manuscript. All authors have read and approved the final version of the manuscript.

## Data Availability

All relevant data have been provided in the main article.

All the raw data related to this study will be deposited into the Repositorio de Datos Académicos UNR (RDA UNR) https://dataverse.unr.edu.ar/ the institutional data repository of the Universidad Nacional de Rosario. The RDA UNR is an open access platform for dissemination and archiving of university research data that complies with the FAIR principles and is developed under the Dataverse Project technology. In addition, the data will be identifiable via DOI number (https://doi.org/10.57715/UNR/5XIFXN) as persistent unique identifier assigned by RDA UNR.

## Abbreviations

ANOVA: (analysis of variance)
Atx1: (copper chaperone Atx1)
BCS: (bathocuproine disulfonate)
Ccs1: (copper chaperone for SOD1)
COX: (cytochrome *c* oxidase- type *aa*3)
COX: (cytochrome *c* oxidase-type *aa*3)
CTR1: (copper transporter 1)
Cu: (copper)
DEAE: (diethylaminoethyl)
DMEM: (Dulbecco’s modified Eagle medium)
DMSO: (dimethyl sulfoxide)
DMT-1: (divalent metal transporter 1)
ER: (endoplasmic reticulum)
FBS: (fetal bovine serum)
Fre: (ferric reductase)
Frt1: (fructose transporter)
GSH: (glutathione)
ICP-OES: (inductively coupled plasma-optical emission spectroscopy)
LIT: (liver infusion tryptose)
Mir1: (mitochondrial import receptor mitochondrial phosphate carrier)
MOI: (multiplicity of infection)
MTT: (3-(4,5-dimethylthiazol-2-yl)-2,5-diphenyltetrazolium bromide)
NeoCup: (neocuproine)
NBT: (nitro-blue tetrazolium chloride)
qRT-PCR: quantitative real-time PCR
*Tc*CuATPase: (*Trypanosoma cruzi* copper P-type ATPase)
*Tc*Cox17: (*Trypanosoma cruzi* cytochrome *c* oxidase copper chaperone COX17)
*Tc*Cox11: (*Trypanosoma cruzi* cytochrome *c* oxidase copper chaperone COX11)
*Tc*Cox19: (*Trypanosoma cruzi* cytochrome *c* oxidase assembly factor COX19)
*Tc*Fet3: (*Trypanosoma cruzi* Fe(II) transport multicopper oxidase FET3)
*Tc*FR: (*Trypanosoma cruzi* ferric reductase)
TGN: (trans Golgi network)
*Tc*IT: (*Trypanosoma cruzi* iron transporter)
*Tc*Pic2: (*Trypanosoma cruzi* Pi/Cu carrier isoform 2)
*Tc*Sco: (*Trypanosoma cruzi* synthesis of cytochrome *c* oxidase chaperone)
ROS: (reactive oxygen species)
TrySH: (trypanothione).

## Supporting Information

Table S1. Primers used for RT-qPCR quantification.

