## supporting file for "Copper trafficking in *Trypanosoma cruzi*: the transcriptional response of candidates to balance toxicity and recruitment"

**Table S1. Primers used for RT-qPCR quantification.**

Gene accession numbers that belong to *T. cruzi* Dm28c strain 2017 from TriTrypDB.

| Protein | gene accession number | primers |
|---|---|---|
| *Tc*FR | BCY84_13742, BCY84_13743, BCY84_05177, BCY84_05181, BCY84_05183, BCY84_05184 | Fw_qTcFR CTGTCGTTGCTCATGTTTCC<br>Rv_qTcFR TTTGTGGACCCAGATCGAAG |
| *Tc*IT | BCY84_01266, BCY84_15905 | Fw_qTcIT AGCGCGAAACGTATCATTGC<br>Rv_qTcIT CCGCGTCTGGGAATCATTTG |
| *Tc*CuATPase | BCY84_03133, BCY84_03134, BCY84_03135 | Fw_qTcCuATPase CGGATGTGGATGAACAGATGG<br>Rv_qTcCuATPase GGCGCAAAATCTGAGACAAC |
| *Tc*Fet3 | BCY84_15293 | Fw_qTcFet3 TTTCCACGTTGCTGCCATTG<br>Rv_qTcFet3 TCTTCACATACCGCCACCAC |
| *Tc*Ubiquitin (housekeeping) | BCY84_18965 | Fw_qTcUbiquitin AGGGCATTCCGGGAAAGATG<br>Rv_qTcUbiquitin CCACCACCATGTGCAGAGTT |
